# Impact of combinatorial histone modifications on acetyllysine recognition by the ATAD2 and ATAD2B bromodomains

**DOI:** 10.1101/2022.11.14.516501

**Authors:** Margaret Phillips, Kiera L. Malone, Brian W. Boyle, Cameron Montgomery, Isabelle A. Kressy, Faith M. Joseph, Kathleen M. Bright, Samuel P. Boyson, Sunsik Chang, Jay C. Nix, Nicolas L. Young, Victoria Jeffers, Seth E. Frietze, Karen C. Glass

**Affiliations:** Department of Pharmacology, Larner College of Medicine, University of Vermont, Burlington, VT, 05405, USA; Department of Pharmaceutical Sciences, Albany College of Pharmacy and Health Sciences, Colchester, VT, 05446, USA; Verna & Marrs McLean Department of Biochemistry & Molecular Pharmacology, Baylor College of Medicine, Houston, TX, 77030, USA; Translational Biology and Molecular Medicine Graduate Program, Baylor College of Medicine, Houston, TX, 77030, USA; Department of Biomedical and Health Sciences, University of Vermont, Burlington, VT, 05405, USA; Molecular Biology Consortium, Advanced Light Source, Berkeley, CA, 94720, USA; Department of Molecular, Cellular and Biomedical Sciences, University of New Hampshire, Durham, NH, 03824, USA

## Abstract

The ATPase family AAA+ domain containing 2 (ATAD2) protein, and its paralog ATAD2B, have a C-terminal bromodomain that functions as a ‘reader’ of acetylated lysine residues on histone proteins. Using a structure-function approach, we investigated the ability of the ATAD2 and ATAD2B bromodomains to select acetylated lysine among multiple histone post-translational modifications. Isothermal titration calorimetry experiments revealed that the ATAD2 and ATAD2B bromodomains selectively recognize distinct patterns of acetylated lysine residues on the N-terminal tails of histone proteins. Adjacent methylation or phosphorylation marks were found to either enhance or weaken the recognition of acetylated lysine by the ATAD2/B bromodomains. Complementary structural studies provide mechanistic insights into how residues within the bromodomain binding pocket coordinate the acetyllysine group in the context of adjacent post-translational modifications. Furthermore, we investigated how sequence changes in amino acids of the histone ligands, either as ‘onco’ mutations or as histone variants, impact the recognition of an adjacent acetylated lysine residue. In summary, our study highlights how the interplay between multiple combinations of histone modifications influences the ‘reader’ activity of the ATAD2 and ATAD2B bromodomains, resulting in distinct binding modes of the two bromodomains.

**KEY POINTS:**

- Multiple independent ATAD2 gene duplication events are evident during metazoan evolution, indicating expansion of functionality in the ATAD2 gene family and suggesting distinct functions for ATAD2 and ATAD2B.
- High-resolution structures of the ATAD2 and ATAD2B bromodomains in complex with their histone ligands demonstrate how multiple post-translational modifications are coordinated.
- Recognition of different subsets acetylated histone ligands by the ATAD2 and ATAD2B bromodomains is driven by unique features within the binding pockets of these paralogous proteins.
- Onco-histone mutations and histone variants that change the amino acid sequence of the histone tails modulate the ATAD2 and ATAD2B bromodomain activity.
- This study demonstrates how the combinatorial activity of multiple post-translational modifications forms a histone code and influences the recognition of acetylated lysine by bromodomain-containing proteins.

## INTRODUCTION

DNA is tightly packaged within the eukaryotic nucleus in the form of chromatin. The primary units of chromatin are nucleosomes, comprised of ∼146 base pairs of DNA wrapped around an octamer of histone proteins that includes two copies of each of histone H3, H4, H2A, and H2B (1–3). Histone proteins are subject to various post-translational modifications (PTMs) by the enzymatic addition of covalent chemical groups to their globular domain and to the amino-terminal tails that extrude from the nucleosome core particle. Many histone PTMs, including methylation, phosphorylation, acetylation, ubiquitylation, SUMOylation, glycosylation, and ADP-ribosylation, govern different biological processes by regulating chromatin structure and function (4,5). Distinctive combinations of multiple types of histone PTMs can be present simultaneously on histones, commonly referred to as forming the ‘histone code’ (6). These unique patterns of histone PTMs serve as binding sites for various effector proteins that translate the histone code into biological readouts, including the activation and repression of gene expression (6–8). Despite intensive research to understand the different histone ‘reader’ proteins that selectively recognize individual PTMs, little is known about the combinatorial recognition of different histone modifications.

Among the protein ‘reader’ domains that bind various histone PTMs, bromodomains are acknowledged as the primary reader of acetylated lysine residues (9). Histone acetylation regulates the relaxation of chromatin structure via repulsive electrostatic interactions between the acetyl group and the DNA phosphodiester backbone, permitting the accessibility of sequence-specific transcription factors to the DNA (10,11). Histone acetylation also serves as a docking site for the recruitment and assembly of protein complexes at distinct regulatory elements within the chromatin (12–14). The histone PTM ligand binding specificity of bromodomains is dictated by the amino acid sequence surrounding the acetyllysine modification of the histone protein and by the residues lining the bromodomain binding pocket (15). While high-resolution structures of different bromodomains in combination with acetylated histone ligands have provided critical insight into ligand recognition, the impact of combinatorial patterns of different PTMs on histone ligand selectivity remains uncharacterized.

ATAD2 and ATAD2B are bromodomain-containing proteins that harbor conserved AAA+ ATPase domains (**A**TPases **A**ssociated with diverse cellular **A**ctivities) (16). As paralogs, ATAD2 and ATAD2B share a high degree of amino-acid similarity (97% similar in the ATPase domains and 74% in the bromodomains) (17). Their bromodomains also share high structural similarity and appear to have similar ligand binding preferences (18–21). While ATAD2 has been demonstrated to serve as a chromatin-binding protein and current research indicates it functions in regulating chromatin structure and transcription (19,20), the function of ATAD2B remains elusive.

In this study, we investigated the similarities and differences between the ATAD2 and ATAD2B paralogs. First, we conducted an evolutionary analysis to identify when a gene duplication event occurred to produce these two paralogs. We compared the selectivity of the reader function of the ATAD2 and ATAD2B bromodomains using different histone H4 ligands containing multiple combinations of PTMs, including acetylated lysine adjacent to another acetylation, phosphorylation, or methylation mark. Then, using an X-ray crystallographic structural approach, we solved five novel high-resolution structures of the ATAD2 and ATAD2B bromodomains in complex with multivalent histone H4 ligands. Additionally, using ligands from mutant onco-histones and histone variants we investigated how changes in the histone tail sequence affect the recognition of adjacent acetylated lysine modifications. The results provide new insights into the interaction of the understudied ATAD2B bromodomain with histone H4 PTMs and highlight the potential for combinations of histone modifications to regulate bromodomain selectivity. Our comparative analysis of the ATAD2 and ATAD2B bromodomains reveals their unique binding activities and sheds light on their possible non-redundant biological functions, providing new insights into their evolution and functional divergence.

## MATERIALS AND METHODS

### Evolutionary analysis of ATAD2 proteins

ATAD2 homologues were identified by BLAST searches of type species representing the major clades of the Opisthokonta using the human ATAD2 and ATAD2b protein sequences. Accession numbers for each homologue from NCBI, UniProt or Ensembl databases are as follows: *Saccharomyces cerevisiae,* P40340; *Schizosaccharomyces pombe,* Q9C0W2, O14114; *Monosiga brevicollis,* A9V250; *Amphimedon queenslandica,* XP_011407258.2; *Hydra vulgaris,* XP_002159335.4; *Nematostella vectensis,* XP_032233592.1; *Limulus polyphemus,* XP_013776023.1; *Ixodes scapularis,* XP_029837885.2; *Bombyx mori,* XP_037874642.1; *Biomphalaria glabrata,* XP_013075716.1, XP_013075717.1; *Mya arenaria,* WAR24384.1, XP_052778215.1; *Ciona intestinalis,* XP_026691996.1; *Branchiostoma floridae,* XP_035667749.1; *Leptobrachium leishanense,* A0A8C5MV52, A0A8C5M7M0; *Gallus gallus,* XP_040553931.1, XP_040520727.1; *Lepisosteus oculatus,* XP_015207022.1, XP_015213378.1; *Danio rerio,* E7FE14, E7EXJ5; *Schmidtea mediterranea,* SMED30003605; *Caenorhabditis elegans,* NP_502289.2; *Brugia malayi,* XP_042930281.1; *Strongylocentrotus pupuratus,* XM_030982860, XM_030982840; *Eptatretus burger,* A0A8C4Q726; *Petromyzon marinus,* S4RMC5; *Homo sapiens,* Q9ULI0, Q6PL18. The regions containing the bromodomain sequences were identified in the full-length protein using NCBI Conserved domain database and confirmed by protein alignment. The evolutionary relationship of bromodomain sequences was determined using the Maximum likelihood method and JTT matrix-based model. The tree with the highest log likelihood was selected. Initial trees for the heuristic search were obtained automatically by applying Neighbor-Join and BioNJ algorithms to a matrix of pairwise distances estimated using the JTT model, and then selecting the topology with superior log likelihood value. Evolutionary analyses were conducted in MEGA11 (22).

### Plasmid construction

Human ATAD2 bromodomain-containing protein (residues 966-1112, UniProt code: Q6PL18) was cloned into a pGEX-6P-1 plasmid containing an N-terminal GST tag and a PreScission Protease cleavage site followed by the bromodomain sequence that had been codon optimized by DAPCEL (Cleveland, OH, USA) and synthesized by GenScript (Piscataway, NJ, USA). Human ATAD2B bromodomain-containing protein (residues 953−1085, Uniprot code: Q9ULI0) was a gift from Nicole Burgess-Brown (Addgene plasmid # 39046) was PCR-amplified and cloned into pDEST15 (GlaxoSmithKline) vector containing an N-terminal GST tag and a PreScission Protease cleavage site followed by the bromodomain sequence. Both plasmids (ATAD2/B) were transformed into *Escherichia coli* BL21(DE3) pLysS competent cells (Novagen, MA, USA) for protein expression.

### Protein expression and purification

The GST-ATAD2 bromodomain-containing and GST-ATAD2B bromodomain-containing BL21(DE3) pLysS cells were grown according to our established protocol (21,23). Briefly, the transformed cells were grown at 37 °C in 2 L of Terrific broth (TB) media supplemented with ampicillin and chloramphenicol antibiotics. The cells were grown to an O.D._600 of_ ∼0.6, and then the temperature was reduced to 20°C. The cells were grown to an O.D._600_ between 0.8-1.0 and induced with 0.5 mM and 0.25 mM isopropyl β-D-1-thiogalactopyranoside (IPTG) for GST-ATAD2 and GST-ATAD2B respectively and grown at 20°C overnight. Next, the cells were harvested at 5,000 RPM for 10 min at 4°C. The cell pellet was suspended in 100 mL of lysis buffer containing 50 mM Tris−HCl pH 7.5, 500 mM NaCl, 5% glycerol, 0.05% Nonidet P-40 alternative, 1 mM dithiothreitol (DTT) and supplemented with 0.1 mg/mL of lysozyme, and 1 tablet of protease inhibitor (Pierce protease inhibitor tablets, EDTA-free, Thermo Fisher). The cells were lysed by sonication, and cell lysate was centrifuged at 10,000 RPM for 20 minutes. The supernatant was incubated with glutathione agarose resin (Thermo Scientific) at 4°C with gentle agitation for 1 hour and 30 minutes. The suspension was poured into 25 mL Econo-Column Chromatography Columns (Bio-Rad) and washed with ten times the resin volume of wash buffer (20 mM Tris−HCl, pH 7.5, 500 mM NaCl, 5% glycerol, 1 mM DTT). The GST tag was cleaved off by incubating the washed beads in wash buffer supplemented with PreScission Protease (∼100 mL at 76 mg/mL) (GE Healthcare) overnight at 4°C. The eluted ATAD2/B bromodomain proteins were concentrated to a total volume of approximately 3 mL. The protein concentration was determined using a Pierce BCA protein assay kit (Thermo Scientific) or by measuring the protein absorbance at 280 nM (19,22). The purity of the ATAD2/B bromodomain was confirmed by sodium dodecyl sulfate-polyacrylamide gel electrophoresis (SDS-PAGE) gels stained with GelCode Blue Safe protein stain (Thermo Scientific) (24,25).

### Histone peptide synthesis

All histone H4, H2A.X, and H2A.Z N-terminal tail peptides with and without modifications were purchased from GenScript (Piscataway, NJ, USA). Amino-acid sequence information of each peptide is available in **Table 1**. All H4 peptides were synthesized with N-terminal acetylation (Nα-ac) and C-terminus amidation and were purified by HPLC to 98% purity. Mass Spectrometry was used to confirm their identity.

**Table 1:**
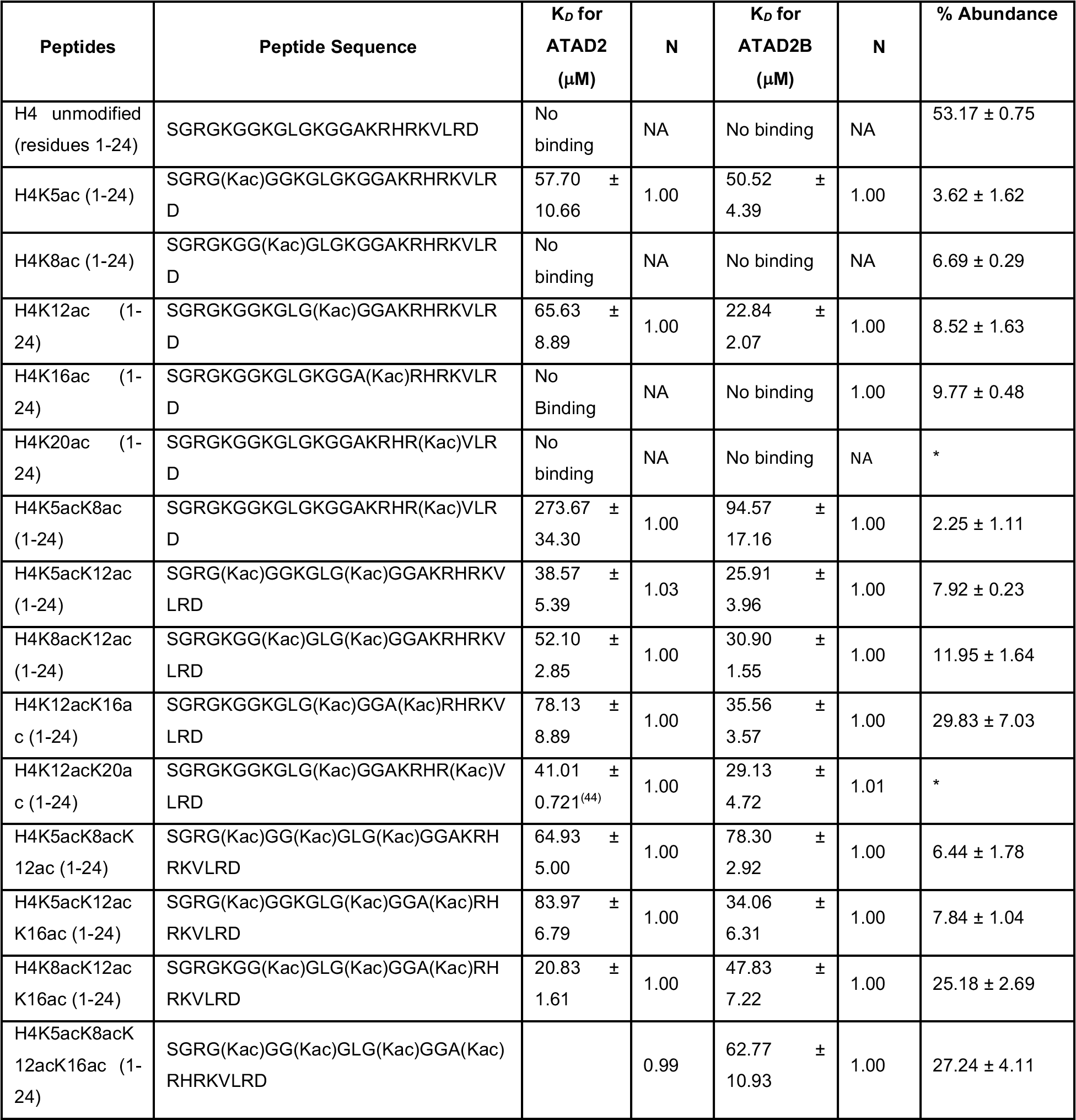
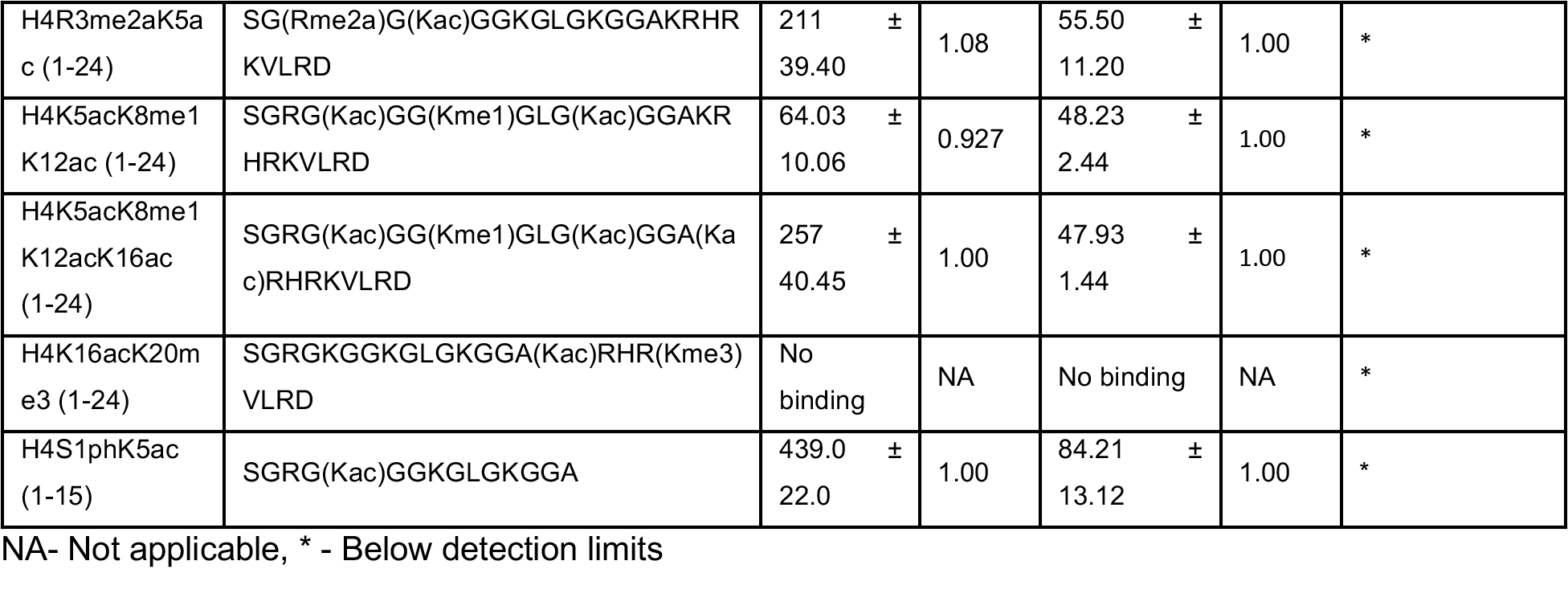
Histone H4 N-terminus tail binding affinities to the ATAD2 and ATAD2B bromodomain-containing proteins. The dissociation constants (*K_D_*) for the ATAD2 and ATAD2B bromodomain interaction with multiple modifications on histone H4 peptides as determined by Isothermal Calorimeter (ITC). The identity and abundance of specific PTMs in MCF7 cells were carried out using mass spectrometry.

### Isothermal titration calorimetry

For isothermal titration calorimetry (ITC), the ATAD2 BRD and the ATAD2B BRD proteins were dialyzed for 48 h into 20 mM sodium phosphate buffer, pH 7.0, 150 mM NaCl, and 1 mM tris(2-carboxyethyl) phosphine (TCEP) (Thermo Scientific) and protein concentration was estimated on a UV-Vis Spectrophotometer at 280 nm. The ITC experiments were carried out at 5°C using a MicroCal iTC200 or a MicroCal PEAQ-ITC instrument (Malvern Analytical). The calorimetric titrations were performed with each histone tail ligand in the syringe (at concentrations of 1.25 mM - 5 mM) and with either the ATAD2 or ATAD2B bromodomain-containing protein in the sample cell (at concentrations ranging from 0.05 mM to 0.2 mM). The histone ligand was added to the sample cell through a series of 20 individual 2 μL injections spaced at time intervals of 150 s while maintaining a continuous stirring speed of 750 RPM. These were preceded by a 0.4 μL preliminary injection of the peptide ligand, excluded from the data integration and binding constant calculation. Control runs of histone peptide into the protein buffer were conducted under identical conditions and subtracted from the ligand into protein runs to correct for the heat of dilution for each peptide. MicroCal PEAQ-ITC software was used for data analysis. All data were fitted using a single set of binding site models to calculate the stoichiometry (N) and the binding constant (*K*_D_). All experiments where binding occurred were performed in triplicate, while runs without binding were performed in duplicate. To calculate the *K*_D_, the average of the three runs was taken as the mean *K*_D_, and the standard deviation was calculated from the mean. All *K*_D_ values and standard deviations for each ligand tested are reported in **Table 1**.

### X-ray crystallography and structure determination

The ATAD2 and ATAD2B bromodomains were further purified for all crystallization experiments through size exclusion chromatography using a Superdex 75 16/60 Sephacryl S-100 high-resolution gel filtration column (GE/Cytiva) on an ÄktaPrime system (GE/Cytiva) in buffer containing 25 mM N-(2-hydroxyethyl)- piperazine-N′-ethane sulfonic acid (HEPES) at pH 7.5, 150 mM NaCl, and 1 mM DTT, and concentrated to 15 mg/mL with Amicon® Ultra (MILLIPORE) concentrators with a 3 kDa MW cut-off (3 MWCO). Protein concentration was estimated using a UV-Vis spectrophotometer at A_280_ nm.

The BRD-histone ligand complex was prepared in a 1:10 molar ratio with the bromodomain at 10 mg/mL and histone ligand at 10 times the molar concentration. For the ATAD2B-H4K12ac (4–17) complex, sitting-drop trays were set up using the Hampton Research Index HT screen in a 96-well VDX plate (Hampton Research, Aliso Viejo, CA, USA) at 4°C. Each 2 μL drop contained 0.5 μL of the complex and 1.5 μL of the mother liquor with a 25 μL reservoir volume. Cryoprotection was achieved by dragging the crystal through LV oil added to the drop. The crystal used for the structure solution grew in 1.8 M ammonium citrate tribasic at pH 7.0 and was mounted on B1A (ALS style) reusable goniometer bases inserted with a 75 μm Dual Thickness Microloop LD built on 18 mm / SPINE length rods (pins) from MiTeGen (Ithaca, NY, USA).

For the ATAD2B-H4S1phK5ac (1–15) complex, sitting drop trays were set up using the Hampton Research Index HT screen in a 96-well VDX plate at 4°C. The 2 μL crystal drop contained 0.5 μL of the complex and 1.5 μL of the mother liquor with a 25 μL reservoir volume. The crystal was harvested using a 300 μm Dual Thickness Microloop LD built on an 18 mm / SPINE length rod (pin) with a B1A (ALS style) reusable goniometer base from MiTeGen. The final crystal was obtained from a condition containing 0.2 M ammonium sulfate, 0.1 M BIS-TRIS (pH 5.5), 25% w/v PEG 3350, and was cryoprotected by sweeping the crystal in the loop through LV oil.

For the ATAD2 BRD H4S1phK5ac (1–15) complex, the ATAD2 BRD protein (10 mg/mL, 0.573 mM) was incubated with the histone peptide (5.73 mM) on ice at a 1:10 molar protein:peptide ratio. Crystallization screens were set up using the Hampton Research Index HT screen in a 96-well, sitting drop VDX plate at 4°C. Each drop (2 µL) consisted of 1 µL of the protein-peptide complex and 1 µL of the mother liquor solution, with a 25 µL reservoir volume. Crystals grew in condition 29 of the Hampton Index HT screen (60% v/v Tascimate^TM^ pH 7.0). The crystal was harvested in a 75 µm Dual Thickness Microloop LD built on 18 mm/SPINE length rods (pins) from MiTeGen (Ithaca, NY, USA). The crystal was cryoprotected with a sweep through 5 µL of LV oil and then was flash frozen in liquid nitrogen.

For the ATAD2B-H4S1CK5ac (1–15) complex, a sitting drop 96 well tray was set up using the Hampton Research Index HT screen, in a 96-well VDX plate at 4°C. The crystal used for structure determination grew in 0.2 M Ammonium sulfate, 0.1 M Tris pH 8.5, 25% w/v PEG 3350. Each drop was 2 μL volume made from 0.5 μL of the complex and 1.5 μL of the mother liquor with a 25 μL reservoir volume. The crystal was harvested using a 200 μm Dual Thickness Microloop LD built on an 18 mm / SPINE length rod (pin) inserted in a B1A (ALS style) reusable goniometer base from MiTeGen. An LV oil sweep was used for cryoprotection.

For the ATAD2 BRD H4S1CK5ac (1–15) complex, the ATAD2 BRD protein (10 mg/mL, 0.573 mM) was incubated with the histone peptide (5.73 mM) on ice at a 1:10 molar protein:peptide ratio. Crystallization screens were set up using the Molecular Dimensions JCSG-plus^TM^ screen and Hampton Index HT screen in a 96-well, sitting drop VDX plate at 4°C. Each drop (2 µL) consisted of 1 µL of the protein-peptide complex and 1 µL of the mother liquor solution, with a 25 µL reservoir volume. Small crystals grew in multiple conditions, but none were satisfactory for collection. Therefore, we set up a separate crystal screen based off a previously successful condition (44). Briefly, we modeled the screen after the Hampton Research Index HT screen conditions 17-18 (1.26 M Sodium phosphate monobasic monohydrate, 0.14 M potassium phosphate dibasic, pH 5., and 0.49 M Sodium phosphate monobasic monohydrate, 0.91 M Potassium phosphate dibasic, pH 6.9). We screened against pH in hanging drop 24-well VDX plates (Hampton Research, Aliso Viejo, CA, USA) over 500 µL of mother liquor solution. In each 2 µL drop, there was 1 µL of the protein-peptide solution and 1 µL of the mother liquor. The optimal crystallization condition was 1.42 M Sodium phosphate monobasic monohydrate and 0.976 M Potassium phosphate dibasic pH 6.4. The crystal was harvested in a 50 µm Dual Thickness Microloop LD built on 18 mm/SPINE length rods (pins) from MiTeGen (Ithaca, NY, USA). The crystal was cryoprotected with a sweep through 28% ethylene glycol in mother liquor, and then was flash frozen in liquid nitrogen.

Data were collected at the Advanced Light Source at the Lawrence Berkeley National Lab on beamline 4.2.2, equipped with an RDI CMOS-8M detector. The diffraction data were processed using XDS (26), and the structure was solved by a molecular replacement using PHASER (27). For all structures, the starting model was a structure of the apo ATAD2B BRD, PDB ID: 3LXJ (28). After obtaining an initial solution, several rounds of structure refinement and building were carried out with PHENIX-1.19.2 (29) and COOT 0.8.9.3. A structural model of the ATAD2B bromodomain in complex with the histone ligands was built into the composite omit map, which showed clear density for the ligand using COOT 0.8.9.3 (30), followed by iterative rounds of refinement and rebuilding in PHENIX-1.19.2 (29) and COOT 0.8.9.3. The final structures were deposited into the Protein Data Bank under PDB IDs: 8EOQ, 8ESJ, and 8SDX, respectively, after validation using MolProbity (31). The final structures were solved at 1.85 Å for ATAD2-H4S1phK5ac (1–15) and at 2.01 Å for ATAD2-S1CK5ac (1–15) and were deposited into the Protein Data Bank under PDB IDS: 8SDQ and 8SDO respectively after validation using MolProbity (28). The hydrogen bonds and hydrophobic interactions were determined using the PLIP (32) program.

### Mass Spectrometry and H4 proteoform quantitation

Cell pellets were collected from plates of MCF-7 cells, and the histones were acid extracted after nuclei isolation as described above and previously in (33). Isolated histones were then resuspended in 85 μL 5% acetonitrile and 0.2% trifluoroacetic acid (TFA) to prepare for high-performance liquid chromatography (HPLC) separation. The histones were separated by type into families and specific variants as described previously (Holt et al., 2021). Briefly, Reverse Phase HPLC fractionation was performed with a Thermo U3000 HPLC system (Thermo Fisher Scientific, Waltham, MA) with a 150 × 2.1–mm Vydac 218TP 3-μm C18 column (HiChrom, Reading, UK, part. no. 218TP3215), using a linear gradient at a flowrate of 0.2 mL/min from 25% B to 60% B for 60 min. The composition of buffers used were A: 5% acetonitrile and 0.2% TFA; B: 95% acetonitrile and 0.188% TFA. After chromatographic separation and histone fraction collection, histone H4 was selected for further study, dried down and resuspended in additional MS buffer A (2% acetonitrile, 0.1% formic acid) for mass spectrometric analysis.

Dried H4 fractions were then diluted using the following calculation, μg H4 = (Peak area –6.0114)/31.215 to calculate the dilution to 200 ng H4/μL in MS buffer A (2% acetonitrile, 0.1% formic acid). 1 μL (200 ng) histone H4 was then loaded onto a 10 cm, 100 μm inner diameter C3 column (ZORBAX 300SB-C3 300 Å 5 μm) for online high-performance liquid chromatography (HPLC) on a Thermo U3000 RSLC nano Pro-flow system. A 70-minute linear gradient using buffer A: 2% acetonitrile, 0.1% formic acid, and B: 98% acetonitrile and 0.1% formic acid was used, and the samples were maintained at a temperature of 4°C. The column eluant was introduced into a Thermo Scientific Orbitrap Fusion Lumos by nanoelectrospray ionization. A static spray voltage of 1800 V and an ion transfer tube temperature of 320°C were set for the source. The MS1 experiment used a 60-k resolution setting in positive mode. An AGC target of 5.0e5 with 200 ms maximum injection time, three micro scans, and a scan range of 700-1400 m/z were used. A targeted charge state of +15 was selected for histone H4. An intensity threshold of 1e5 was set for the selection of ions for fragmentation. The target precursor selected for MS2 fragmentation included all major H4 peaks and chose the top 20 most abundant m/z. ETD fragmentation at a 14 ms reaction time, 5.0e5 reagent target with 200 ms injection time was used. MS2 acquisition was performed using the orbitrap with the 60 k resolution setting, an AGC target of 5.0e5, a max injection time of 200 ms, a ‘normal’ scan range, and three micro scans.

Identification of H4 at the desired charge state of +15 was observed in mass spectra. Raw files were converted to mzXML. Data processing was performed by a custom analysis suite as previously described (33,34).

## RESULTS

### Gene duplication of *ATAD2* occurred several times during eukaryotic evolution

The *ATAD2* gene encodes for two AAA+ ATPase domains and a C-terminal bromodomain that is thought to play an essential role in chromatin remodeling. *ATAD2* is found in a diverse variety of organisms ranging from yeast to humans. A gene duplication event resulting in two paralogous proteins, ATAD2 and ATAD2B, found in humans and many other higher eukaryotic organisms. However, the origin and conservation of ATAD2/B is unclear.

Evolutionary analysis of ATAD2 homologs across eukaryotes identifies at least a single homolog of ATAD2 in all species within the opisthokonts, the major eukaryote super-group representing fungi and metazoans (**Figure 1A**). Within the vertebrates, most species contain both ATAD2 and ATAD2B homologs, with the exception of the early-branching chordates, suggesting that the gene duplication occurred more recently during vertebrate evolution. One of the early-branching chordates, the purple sea urchin (*Strongylocentrotus purpuratus*), also contains two ATAD2 homologues. Further comparison of the protein sequences of the purple sea urchin homologues with ATAD2 and ATAD2B sequences from other vertebrates reveals that these are more divergent from the mammalian ATAD2 proteins. This comparison also indicates a more recent expansion of ATAD2 in the Chordata after divergence from the evolutionary trajectory of mammals.

**Figure 1.**
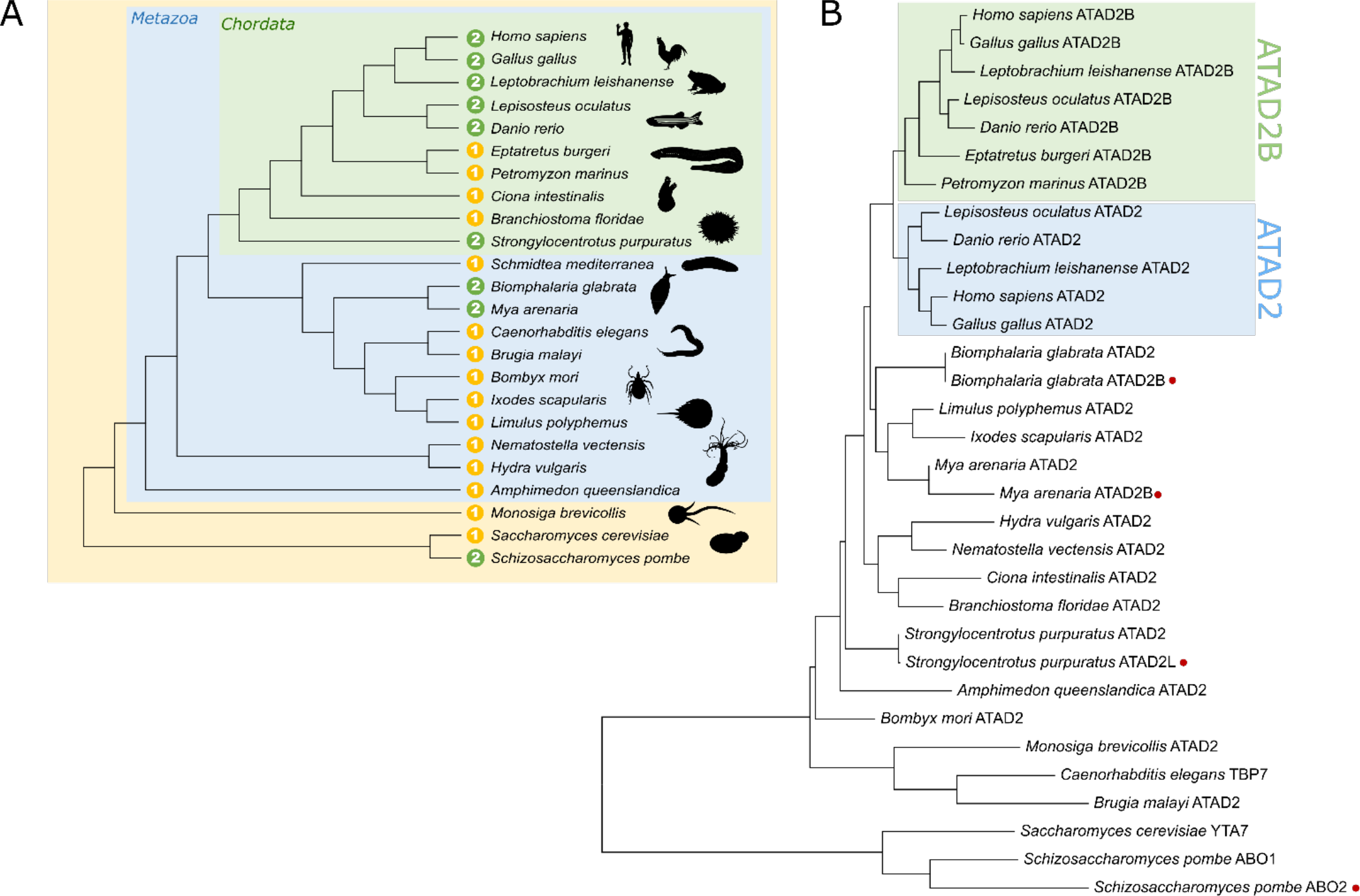
Evolution of ATAD2 homologues in animals. (A) Phylogeny of species within the Opisthokonta with the number of ATAD2 homologues indicated in the yellow (one homologue) or green (two homologues) circles. (B) Phylogenetic tree of the bromodomain sequences in ATAD2 homologues from species depicted in A, inferred by using the maximum likelihood method and JTT matrix-based model. The branches representing the ATAD2 and ATAD2B lineages in vertebrates are highlighted in the blue and green shaded boxes respectively. Red circles indicate predicted ATAD2 duplications that occurred independently of vertebrate ATAD2 and ATAD2B evolution. The tree is drawn to scale, with branch lengths measured in the number of substitutions per site. Animal silhouette images were obtained from www.phylopic.org.

To determine how the evolution of ATAD2 orthologues might have functional implications for the bromodomain, we performed a phylogenetic analysis of ATAD2 bromodomain sequences from the selected species within the Opisthokonta. ATAD2 and ATAD2B bromodomains assemble into two distinct groups in the vertebrates, supporting the hypothesis that the second paralog derived from a single gene duplication event. Interestingly, the single ATAD2 homologues we identified in jawless fish (*E. burgeri* and *P.* marinus) group with the ATAD2B homologs from other species (**Figure 1B**, green shaded box). This presents two possibilities that either ATAD2B was the original ATAD2 in vertebrates that was later duplicated to form ATAD2 after the divergence of the jawless fish, or that the ATAD2 homolog was lost at some point during the jawless fish evolutionary trajectory. This analysis also reveals evidence of multiple independent duplications of ATAD2 in many of the other species we examined (*M. arenaria*, *B. glabrata* and *Schiz. pombe*), suggesting that expansion of ATAD2 variants is a relatively common evolutionary event and might represent a beneficial strategy for dealing with increasing transcriptional complexity.

Gene duplication and retention of the ATAD2 orthologs in vertebrates suggest that the ATAD2 and ATAD2B proteins may have independent functions. As paralogs, human ATAD2 and ATAD2B share a high degree of amino-acid similarity (97% similarity in the ATPase domains and 74% in the bromodomains) (17). Their bromodomains also share high structural similarity and have similar ligand binding preferences. Their conserved structural features make it difficult to determine if ATAD2 and ATAD2B have independent or redundant functions. The bromodomains of ATAD2 and ATAD2B have been shown to preferentially recognized histone H4K5acK12ac (19,21,23,35), a post-translational modification associated with newly synthesized histones. ATAD2 has been demonstrated to serve as a chromatin-binding protein, and current research indicates it regulates chromatin structure and transcription (19,20). Expression of ATAD2 is restricted to S-phase (19), indicating a cell cycle-specific function of the ATAD2 complex. However, the cellular function(s) of ATAD2B is still unknown. Here we conducted an in-depth structure and function analysis of the ATAD2 and ATAD2B bromodomains to characterize further their unique bromodomain features that contribute to distinct histone ligand recognition and binding specificities.

### Combinatorial histone PTMs influence ‘Kac’ recognition by the ATAD2/B bromodomains

Our previous study compared the recognition of acetylated histone ligands by the ATAD2 and ATAD2B bromodomains using an unbiased histone PTM ligand binding dCypher screen that included 288 distinct peptides with both single and combinatorial histone modifications (21). We found that the ATAD2 bromodomain preferred a select subset of acetyllysine modifications, recognizing 11 histone ligands within the context of adjacent acetylation, phosphorylation, and methylation modifications (21). In contrast, the ATAD2B bromodomain appeared less selective and could bind acetylated lysine with limited influence from adjacent PTMs (21). While the bromodomains recognize only acetylation marks, we hypothesized that neighboring PTMs (of varying shape, size, and charge) would impact the acetyllysine insertion into the bromodomain binding pocket.

Thus, to understand how adjacent PTMs modulate the binding activities of the ATAD2 and ATAD2B bromodomains, we first performed a quantitative analysis (using ITC) of the binding of these two bromodomains with histone H4 peptides containing combinatorial PTMs. The histone peptides used in the experiments were selected from the top histone H4 ligands identified previously(21) and were designed to have uniform amino-acid sequence lengths to allow for a direct comparison (**Table 1**, **Supp. Fig. S1 and S2**).

Among the mono-acetylated histone H4 ligands tested, the ATAD2 bromodomain prefers the H4K5ac modification binding with an affinity of *K_D_* = 57.70 ± 10.66 μM compared to the ATAD2B bromodomain, which demonstrates the highest binding affinity for the H4K12ac ligand (*K_D_* = 22.84 ± 2.07 μM). Neither ATAD2/B bromodomains bind to H4K8ac, H4K16ac, and H4K20ac peptides.

Among the di-acetylated histone H4 peptides tested, both ATAD2/B bromodomains prefer the H4K5acK12ac modification, binding with affinities of *K_D_* = 38.57 ± 5.39 μM and 25.91 ± 3.96 μM, respectively. In contrast, significantly reduced binding affinities are observed for recognition of the H4K5acK8ac combinatorial marks by the ATAD2/B bromodomains (*K_D_* = 273.67 ± 34.30 μM and 94.57 ± 17.16 μM, respectively), when compared to H4K5ac alone.

For the three tri-acetylated H4 peptides tested, the ATAD2 bromodomain binds to H4K8acK12acK16ac peptide with the highest affinity (*K_D_* = 20.83 ± 1.61μM) but displays an almost four-fold decrease in affinity for H4K5acK12acK16ac (*K_D_* = 83.97 ± 6.79 μM). In contrast, the ATAD2B bromodomain shows the tightest binding to the H4K5acK12acK16ac combinatorial modifications (*K_D_* = 34.06 ± 6.31 μM).

The differences in ATAD2 and ATAD2B bromodomain binding selectivity are most noticeable in H4 peptides carrying hydrophobic methyl modifications in combination with acetylation marks. Overall, the binding affinity of the ATAD2 bromodomain for acetyllysine marks near mono- and di-methylation modifications is reduced, while the presence of adjacent methyl groups has an insignificant effect on the ATAD2B bromodomain activity (**Table 1**). For example, the addition of bulky di-methyl groups at Arg 3 significantly reduces the affinity of the ATAD2 bromodomain for the ‘K5ac’ mark in H4R3me2aK5ac (*K_D_* = 211 ± 39.40 μM compared to the H4K5ac *K_D_* = 57.70 ± 10.66 μM). However, the ATAD2B bromodomain tolerates the presence of adjacent methyl groups, with no notable change in its affinity for the H4R3me2aK5ac (*K_D_* = 55.50 ± 11.20 μM) compared to H4K5ac alone (*K_D_* = 50.52 ± 4.39 μM).

Histone post-translational modifications are dynamic and often found in different abundances on the N- and C-terminal regions of histone proteins (6,36–40). The modified histone ligands in the dCypher screen were designed and synthesized to cover known post-translational modifications on the canonical and variant histones in a combinatorial manner. However, it does not necessarily represent the physiologically relevant combinations of modifications on individual histone molecules (also known as histone proteoforms) that are biologically available within the cell. To assess this, we used a top-down proteomic approach, which omits protein digestion, to identify and quantify the abundance of intact histone H4 proteoforms in MCF7 human breast cancer cells (**Table 1**, right column). This proteoform profile confirms that mono-, di-, tri-, and tetra-acetylated histone H4 are biologically relevant ligands and that phosphorylation and methylation marks surrounding the acetylated lysine modifications are also available within these cells. Thus, the combination(s) of physiologically relevant combinations of histone PTMs likely influence the histone recognition activity of the ATAD2 and ATAD2B bromodomains within the cell.

### Molecular basis of acetyllysine recognition by the ATAD2/B bromodomains

#### (A)#Distinct Recognition of mono-acetylated histone H4

While recognition of mono-acetylated histone ligands such as H4K5ac and H4K12ac by the ATAD2 bromodomain is established (20,23), there is little information on how its paralog, the ATAD2B bromodomain, coordinates histone H4 acetylated lysine modifications. To elucidate the molecular mechanisms involved in acetyllysine recognition by the lesser-studied ATAD2B, we co-crystallized the ATAD2B bromodomain (BRD) with the mono-acetylated histone H4K12ac (residues 4-17) ligand. The X-ray crystal structure of this complex was solved to a resolution of 2.0 Å by molecular replacement using the apo ATAD2B-BRD crystal structure (PDB ID: 3LXJ) as the search model (**Figure 2A, Supp. Table S1**). Good ligand density is observed for residues Gly 6 to Arg 17 of the histone H4 peptide within the canonical ATAD2B bromodomain pocket (**Figure 2B)**. Multiple hydrogen bonds and hydrophobic interactions contribute to the histone H4K12ac ligand coordination (**Figure 2C**). While the ATAD2 and ATAD2B bromodomains share a conserved acetyllysine recognition pattern, the two proteins’ coordination of the surrounding histone residues by the bromodomain binding pocket is starkly different. A detailed comparison of the ATAD2 BRD in complex with H4K12ac ligand solved previously (PDB ID: 4QUT) (20) (shown in **Figure 2D**), with our ATAD2B BRD complex structure highlights some of the features of coordinating H4K12ac by the two bromodomains (compare **Figure 2C** and **D)**. Coordination of the acetyllysine moiety carbonyl group occurs via a direct hydrogen bond to the conserved N1038 and N1064 of the ATAD2B and ATAD2 bromodomains, respectively. Similarly, the hydrophobic interaction between V987 and the gatekeeper residue I1048 of ATAD2B with the acetyllysine group of the histone ligand is similar to the hydrophobic interaction made by the corresponding V1013 and I1074 residues in the ATAD2 bromodomain. However, major differences in the coordination of the histone H4K12ac ligand by the two bromodomains lie in the extent of the histone backbone readout. In ATAD2B, multiple hydrogen bonds directly coordinate the histone backbone within the ZA/BC loop region of the bromodomain, while only three (one through water) hydrogen bonds are observed within the ATAD2 BRD pocket. In our ATAD2B structure, the conserved tyrosine Y1037 makes direct hydrogen bonds with the histone ligand’s Gly 9 and Gly 11. In contrast, in the ATAD2 structure (4QUT), there is only one water-mediated bond to Gly 11 of the histone ligand through the backbone carbonyl group of the conserved tyrosine Y1063. Furthermore, we observe multiple hydrogen bonds between the histone ligand backbone and ATAD2B bromodomain residues. These include Gly 7 of histone H4 and L1035 and R1049 of ATAD2B, Lys 8 and Gly 9 with E1036, Leu 10 with D994, and Arg 17 with S993, for a total of 9 direct hydrogen bonds coordinating the ATAD2B BRD – H4K12ac ligand interaction in addition to contacts with the acetyllysine group. Thus, our crystal structure illustrates how the ATAD2B BRD uses a canonical binding pocket to ‘read’ acetylated histone H4K12. It also highlights the diverse network of polar and non-polar interactions that guide the selection of this modification. Importantly, despite sharing high structural similarity in recognition of acetylated lysine, the ATAD2 and ATAD2B bromodomains coordinate the histone H4 protein using a distinct binding mode that is fine-tuned through specific interactions between the bromodomain binding pockets and the histone H4 backbone residues.

**Figure 2:**
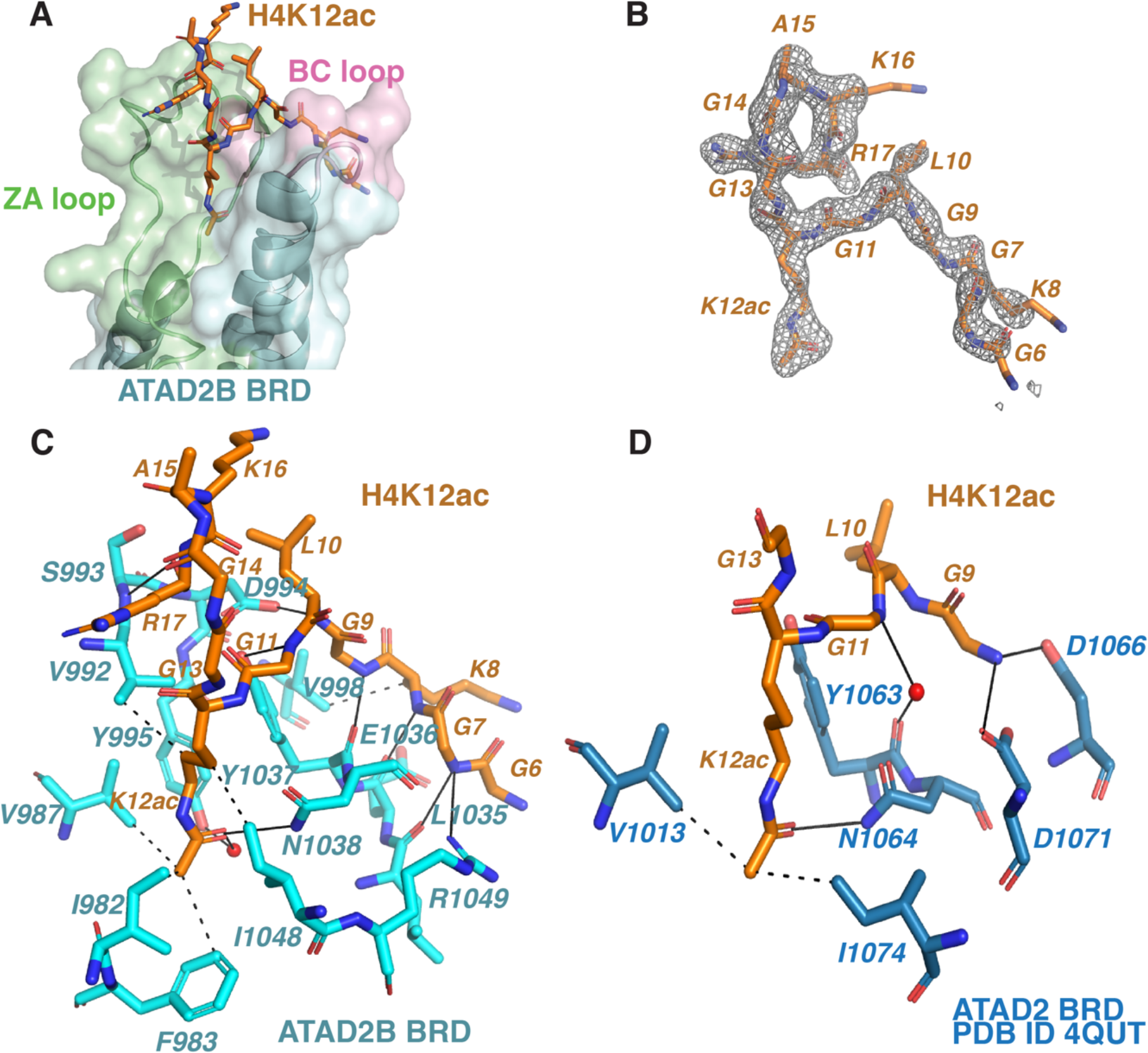
Coordination of the histone H4K12ac ligand by the ATAD2/B bromodomains. (A) Surface representation of the ATAD2B BRD in complex with the H4K12ac ligand (residues 4-17) (PDB ID: 8EOQ). The H4K12ac ligand residues are shown as orange sticks, while the cartoon representation of BRD (cyan) highlights the ZA and BC loops in green and pink, respectively. (B) An isolated image of the simulated annealing composite omit map (gray) around the histone ligand H4K12ac (orange) contoured at 1σ. The composite omit map was calculated before building the histone ligand into the structure. (C) Coordination of the H4K12ac ligand by the ATAD2B BRD. Residues lining the BRD pocket involved in ligand coordination are displayed in cyan, while the H4K12ac ligand residues are in orange. Hydrogen bonds are represented by solid black lines, while dashed black lines show hydrophobic interactions. Water molecules are shown in red. (D) Coordination of the H4K12ac ligand (residues 9-16) by the ATAD2 BRD as obtained from the previously solved crystal structure (PDB ID: 4QUT) (1,2). Residues lining the BRD pocket involved in ligand coordination are displayed in dark blue, while the H4K12ac ligand residues are in orange. Hydrogen bonds are represented by solid black lines, while dashed black lines show hydrophobic interactions. Water molecules are shown in red. All figures were made using the PyMOL Molecular Graphics system using version 2.3, Schrödinger, LLC, and all contacts were determined using the PLIP program (3).

#### **(B)** Effect of adjacent modifications on ‘Kac’ recognition

Based on the ATAD2B-H4K12ac structure, where several polar contacts are observed between histone residues surrounding the acetyllysine mark and the bromodomain, we hypothesized that neighboring PTMs would impact acetyllysine recognition by the ATAD2/B bromodomains. Our previous study highlights the importance of the first three residues, ‘SGR,’ in coordinating acetyllysine on the histone H4 tail as their deletion completely abolished H4K5ac and H4K8ac ligand binding (21). Based on this, we selected the histone H4S1phK5ac (residues 1-15) ligand as we hypothesized that a negatively charged phosphoryl group (ph) at Ser 1 would impact recognition of the adjacent H4K5ac group by the two BRDs. To characterize how this H4S1ph modification alters the molecular mechanism of acetyllysine coordination, we co-crystallized the ATAD2 and ATAD2B bromodomains with the H4S1phK5ac peptide. We solved the crystal structures of the ATAD2 and ATAD2B bromodomain-ligand complexes at resolutions of 1.85 Å (PDB ID: 8SDQ) and 1.4 Å (PDB ID: 8ESJ), respectively (**Figure 3**, and **Supp. Tables S2** and **S3,** respectively). Electron density for histone H4 residues 1-7 is observed within the ATAD2 BRD binding pocket (**Figure 3A, C**), whereas electron density for H4 residues 1-6 is seen in the ATAD2B BRD pocket (**Figure 3B, D**). Similar to H4K12ac, coordination of H4K5ac by both bromodomains involves a combination of hydrogen bonds and hydrophobic contacts. In our ATAD2/B structures, direct hydrogen bonds are observed between conserved N1064 of ATAD2 and corresponding N1038 in ATAD2B and the carbonyl of the H4K5ac group (**Figure 3C, D**). This acetyllysine residue is further stabilized by hydrophobic contacts between its sidechain and residues F1009, V1013, and V1018 of the ATAD2 BRD (**Figure 3C**), and the corresponding residues F983 and V987 in the ATAD2B BRD (**Figure 3D**). The conserved tyrosine residue (Y1063 in ATAD2 and Y1037 in ATAD2B BRD) is also crucial for coordinating the histone H4 ligand as seen by the hydrogen bonds between this tyrosine residue and Gly 2 and Gly 4 of the H4 ligand in the two structures (**Figure 3C, D)**. Similarly, coordination of the adjacent phosphorylated Ser 1 in the histone H4 ligand is comparable between the two bromodomains where the backbone carbonyl of the glutamic acid residue (E1062 in ATAD2 and E1036 in ATAD2B) interacts with the backbone amide of the Ser1ph group. Interestingly, while the ATAD2 BRD coordinates several residues in the histone H4 ligand via water-mediated hydrogen bonds (D1020 with Arg 3 and Gly 2, and D1071 with Gly 4), all of these parallel interactions in the ATAD2B BRD occur through direct hydrogen bonds (**Figure 3D**). Notably, the direct hydrogen bond interactions between the ATAD2B BRD- H4S1phK5ac ligand appear to stabilize its coordination resulting in an increased binding affinity of *K_D_* = 84.21 ± 13.12 μM when compared with the ATAD2 BRD-H4S1phK5ac *K_D_* = 439.0 ± 22.0 μM (**Table 1**). Overall, our structure-function analysis of the ATAD2 and ATAD2B bromodomains demonstrates that in the context of multiple post-translational modifications adjacent to acetyllysine, such as the phosphoryl group, differences in the amino acid residues of the bromodomain binding pockets dictate a unique readout of modified histone H4 by ATAD2 and ATAD2B. The differences in their histone binding modes contribute to the distinct binding specificities observed for the bromodomains and may contribute to different functional activities of the full-length ATAD2 and ATAD2B proteins in the cell.

**Figure 3:**
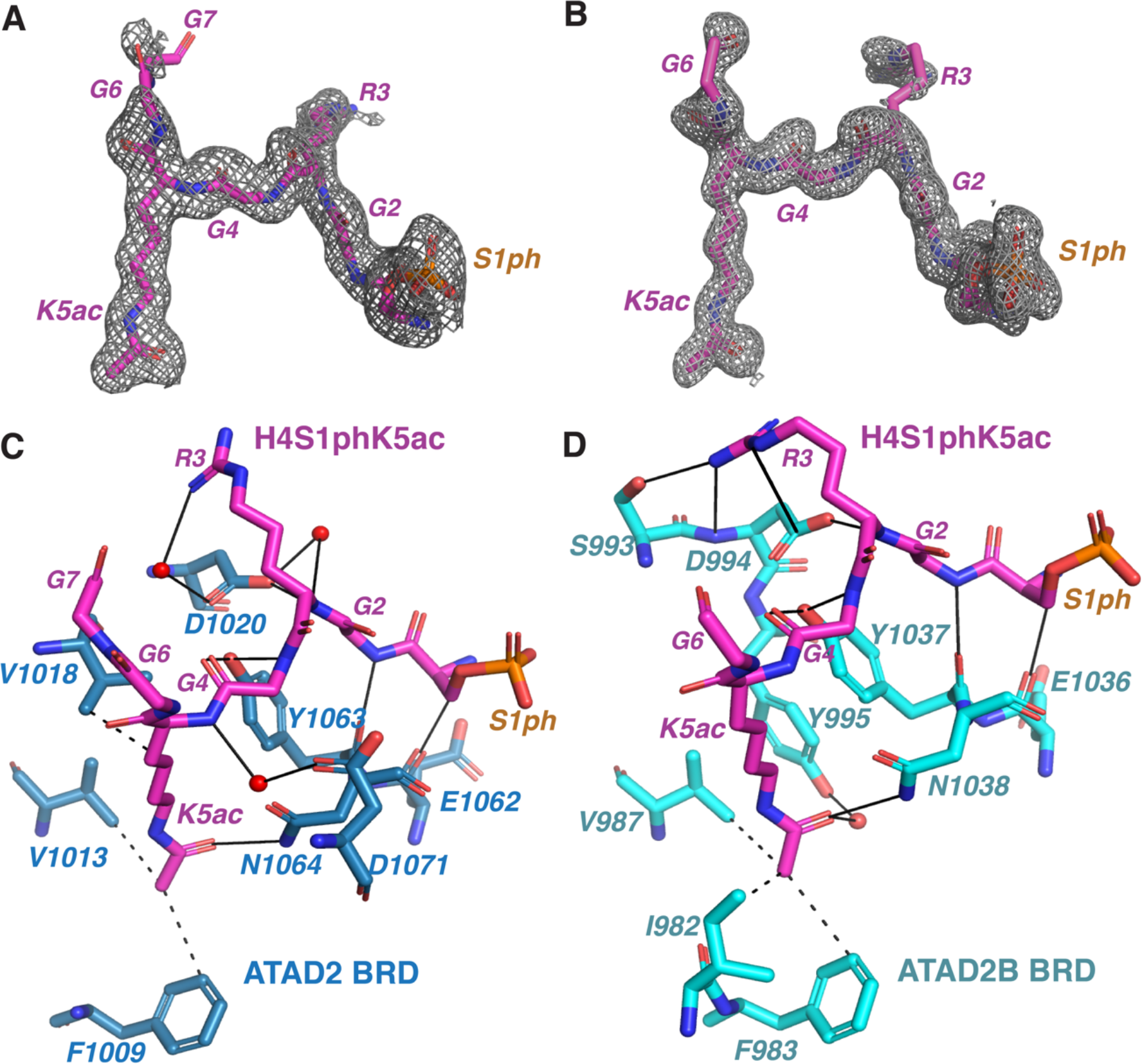
Coordination of the histone H4S1phK5ac ligand by the ATAD2A and ATAD2B bromodomains. (A) An isolated image of the simulated annealing composite omit map (gray) around the histone ligand H4S1phK5ac (magenta, residues 1-7) contoured at 1σ observed in the ATAD2 BRD- H4S1phK5ac complex. The composite omit map was calculated before building the histone ligand into the structure. (B) An isolated image of the simulated annealing composite omit map (gray) around the histone ligand H4S1phK5ac (magenta, residues 1-6) contoured at 1σ observed in the ATAD2B BRD-H4S1phK5ac complex. (C) Coordination of the H4S1phK5ac ligand (residues 1-7) by the ATAD2 BRD. Residues lining the BRD pocket involved in ligand coordination are displayed in dark blue, while the H4S1phK5ac ligand residues are shown in magenta. (D) Coordination of the H4S1phK5ac ligand (residues 1-6) by the ATAD2B BRD. Residues lining the BRD pocket involved in ligand coordination are displayed in cyan, while the H4S1phK5ac ligand residues are shown in magenta. Hydrogen bonds are represented by solid black lines while dashed black lines show hydrophobic interactions. Water molecules are shown in red. All figures were made using the PyMOL Molecular Graphics system using version 2.3, Schrödinger, LLC, and all contacts were determined using the PLIP program (3).

### The Histone Amino Acid Sequence Regulates Bromodomain Activity

The histone backbone residues are crucial in coordinating the ’Kac’ mark by participating in water-mediated or direct polar contacts with the bromodomain binding pocket. We hypothesized that any changes in the sequence of the N-terminal histone tail would impact the recognition of nearby acetylated lysine modifications by the bromodomain.

#### **(A)** Mutations in the H4 N-terminal tail alter the ATAD2/B bromodomain activity

A recent study identified ‘onco’ mutations in the histone genes associated with cancer (41). Several of these mutations occur at or adjacent to sites of known PTMs (41,42). We hypothesized that these ‘onco’ mutations would affect how the ATAD2/B bromodomains ‘read’ the closely associated histone acetylation marks. In this study, we selected the top four N-terminal mutations near the site of Lys 5 acetylation and assessed their impact on ATAD2/B bromodomain binding activity.

Surprisingly our ITC data demonstrate a three-fold increase in binding affinity of the ATAD2 BRD for the onco-histone H4S1C with a K5ac modification when compared to the wild-type histone H4K5ac ligand (23) (**Table 2, Supp. Figure S3**). However, the ATAD2B BRD binding activity decreases in response to the same onco-histone mutation (**Table 2, Supp. Figure S4**) (21). The onco-histone H4R3C mutation also increases the binding affinity of the ATAD2 BRD for H4K5ac but decreases the binding affinity of the ATAD2B BRD for H4K5ac. Interestingly, mutation of histone H4G4 to either Asp or Ser abolishes the recognition of K5ac by both the ATAD2 and ATAD2B bromodomains (**Table 2**).

**Table 2:**
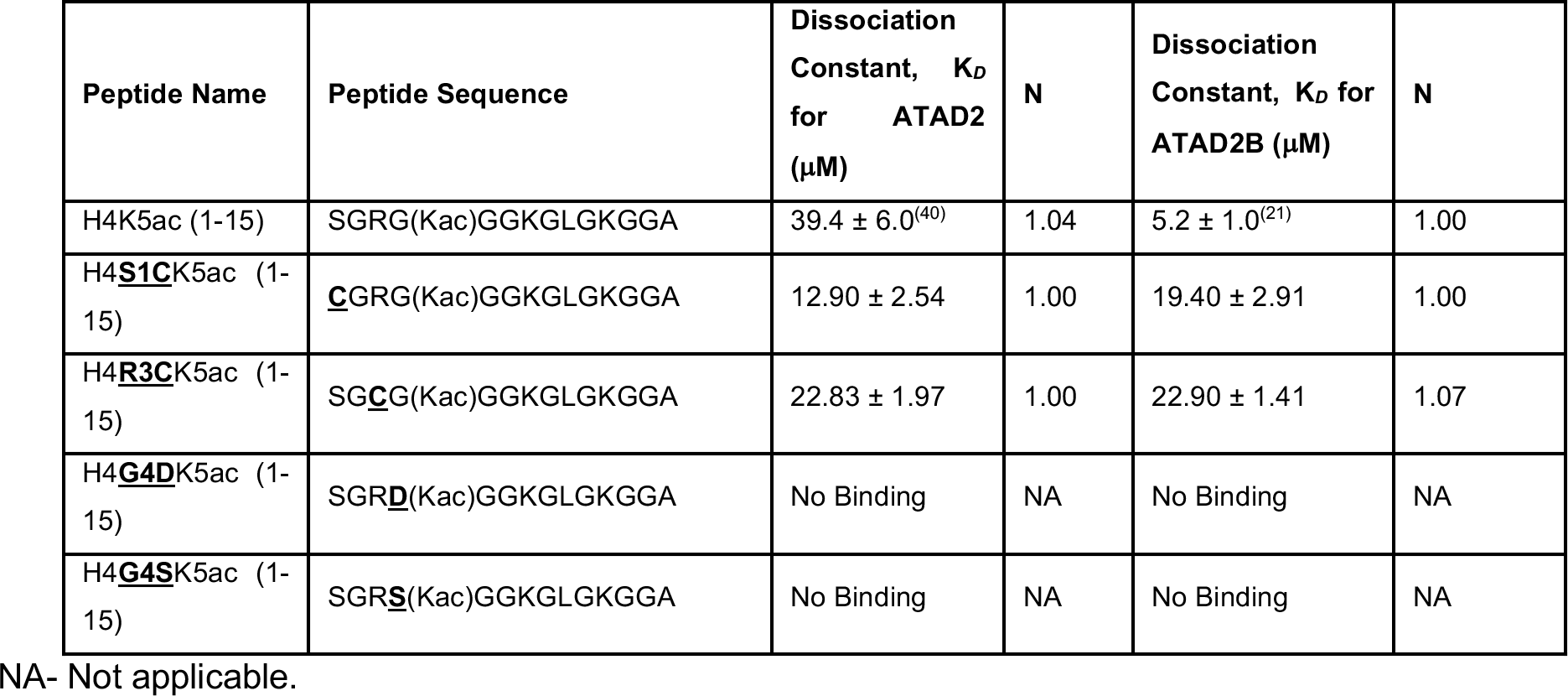
Binding affinities of the ATAD2/B bromodomain with the histone H4 oncohistone peptides with an acetyllysine modification on K5 as measured by ITC. The apparent dissociation constants (*K_D_*) are given in micromolar. Amino acid mutations are highlighted in bold and underlined in the sequence.

To further investigate the molecular mechanisms involved in the acetyllysine recognition of the mutated onco-histone H4 tail, we solved X-ray crystal structures of the ATAD2 and ATAD2B bromodomains in complex with H4S1CK5ac (residues 1-15) at resolutions of 2.01 Å (PDB ID: 8SDO, **Figure 4A, and C, Supp. Table S4**) and 2.79 Å (PDB ID: 8SDX, **Figure 4B and D, Supp. Table S5**), respectively. As shown in **Figures 4A and B**, the 2F_o_–F_c_ composite omit maps show that ligand density is observed for the first six residues in the ATAD2 BRD pocket, and the first seven residues of the ATAD2B BRD pocket. The coordination of the K5ac group of the onco-histone H4S1CK5ac ligand is well conserved between the ATAD2 and ATAD2B bromodomains (**Figures 4C and D**). The higher affinity of the ATAD2 BRD for the S1C mutant compared to the wild-type H4K5ac (**PDB ID: 7M98)(Supp. Figure S6)** (23) can be explained by the increased water-mediated hydrogen bonds observed between Cys 1 of the H4 ligand and the E1062 residue in the ATAD2 BRD, along with an additional hydrophobic contact between the K5ac group and the gatekeeper residue I1074 in ATAD2 BRD.

**Figure 4:**
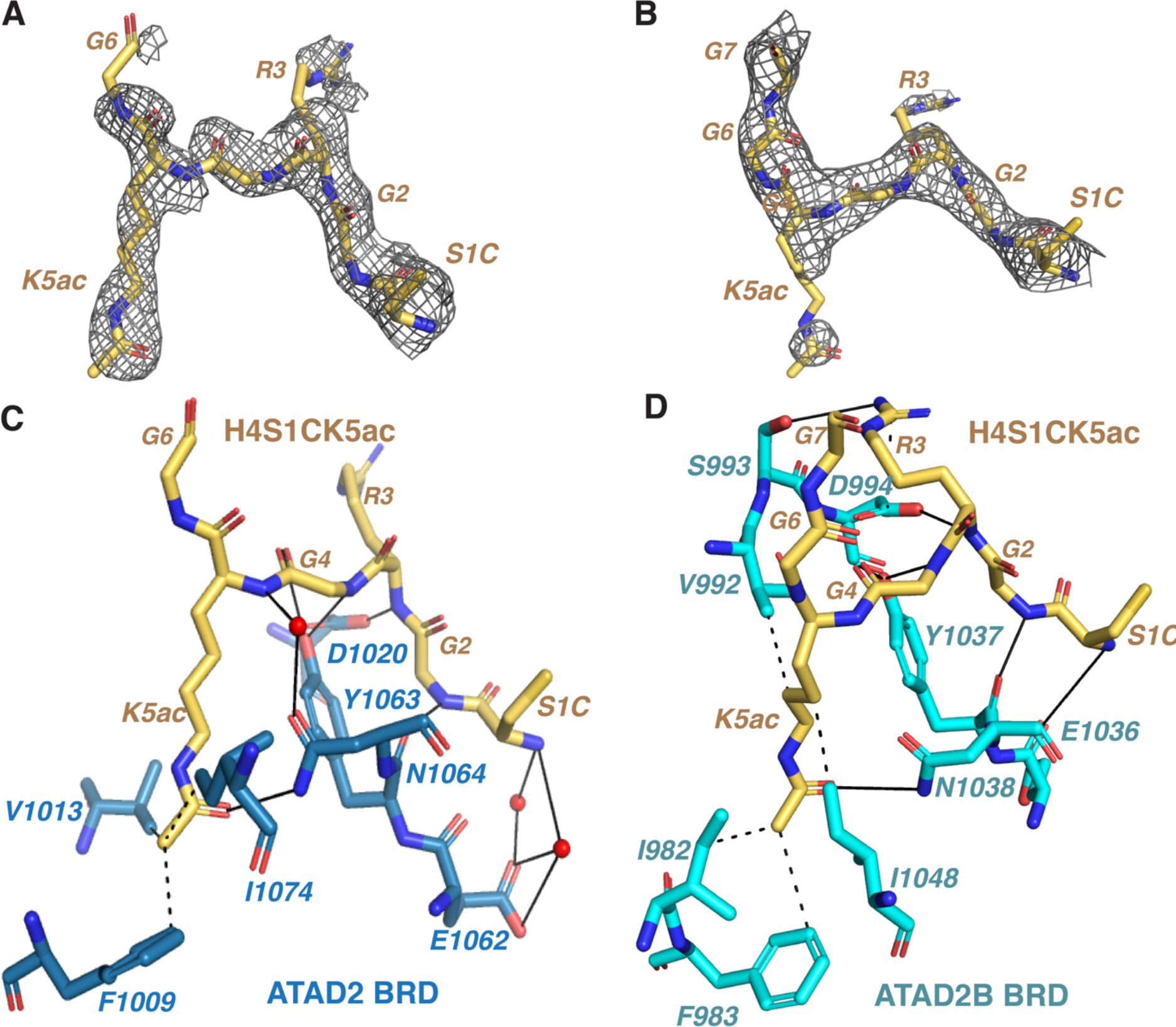
Coordination of the oncohistone H4S1C with K5ac modification by the ATAD2/B bromodomains. (A) An isolated image of the simulated annealing composite omit map (gray) around the oncohistone H4S1C with K5ac (yellow, residues 1-6) contoured at 1σ observed in the ATAD2 BRD- H4S1CK5ac complex. The composite omit map was calculated before building the histone ligand into the structure. (B) An isolated image of the simulated annealing composite omit map (gray) around the histone ligand oncohistone H4S1C with K5ac (yellow, residues 1-7) contoured at 1σ observed in the ATAD2B BRD- H4S1phK5ac complex. (C) Coordination of the H4S1C with K5ac ligand (residues 1-6) by the ATAD2 BRD. Residues lining the BRD pocket involved in ligand coordination are displayed in dark blue, while the H4S1CK5ac ligand residues are shown in yellow. (D) Coordination of the H4S1CK5ac ligand (residues 1- 7) by the ATAD2B BRD. Residues lining the BRD pocket involved in ligand coordination are displayed in cyan, while the H4S1CK5ac ligand residues are shown in yellow. Hydrogen bonds are represented by solid black lines, while dashed black lines show hydrophobic interactions. Water molecules are shown in red. All figures were made using the PyMOL Molecular Graphics system using version 2.3, Schrödinger, LLC, and all contacts were determined using the PLIP program (3).

Due to the lack of an ATAD2B BRD-H4K5ac structure, we cannot directly ascertain how the H4S1C onco-histone mutation affects the K5ac insertion within the ATAD2B binding pocket. However, our structure reveals that K5ac is coordinated by at least four hydrophobic contacts, one of which involves the ATAD2B BRD gatekeeper residue I1048 (**Figure 4D**). Conserved N1038 and Y1037 of ATAD2B BRD form direct hydrogen bonds with the K5ac sidechain and the adjacent Gly 4. The N-terminus of the H4S1CK5ac ligand is stabilized via hydrogen bond formation with the Y1037 and E1036 residues in the ATAD2B BRD. Additional hydrogen bonds are observed between Gly 2 and Arg 3 of the histone H4 ligand and residues D994 and S993 of ATAD2B BRD, respectively.

Comparing the coordination of H4K12ac and H4S1CK5ac by ATAD2B suggests that several hydrogen bond contacts between the BRD binding pocket and the histone backbone may be lost due to this ‘onco’ mutation. Overall, our structural and ITC data demonstrate that ‘onco’ mutations within the histone H4 backbone influence the recognition of adjacent acetyllysine modification and would likely affect the ATAD2/B bromodomain binding activity in a cellular context.

#### **(B)** Histone variants H2A.X and H2A.Z demonstrate distinct ATAD2/B bromodomain activity

Histone variants have unique amino acid sequences compared to canonical histones and play distinct roles in various cellular processes (43). Histone variants differ in the co-translational and post-translational modifications they harbor (44). While many H2A sequences are quite similar, H2A.X and H2A.Z.1 and H2A.Z.2 are more distinct (45). The first five amino acids of H2A.X, SGRGK, are identical to the canonical H2A sequences. Histone H4 also shares an identical starting sequence. This preserves the constitutive N-terminal acetylation and the more abundant K5ac between sequences (44). However, starting at the sixth amino, the sequences diverge slightly H2As (QGGK), H2A.X (TGGK), and H4 (GGK). H2A.Z shares far less homology near common modification sites and is further observed not to be N-terminally acetylated (44,45). Beyond these basic observations, how these variants modulate bromodomain binding activity is understudied. Among the histone variants tested in our previous dCypher screen, ATAD2 and ATAD2B bromodomains preferred the acetylated form of histone H2A.X (21).

To further investigate how changes in histone amino acid sequence influence acetyllysine recognition, we used ITC to calculate the binding affinities of the ATAD2/B bromodomains with histone variants H2A.X and H2A.Z.1 that carry mono- and di- acetylated lysine modifications (**Table 3, Supp. Figure S5**). Compared to binding with the canonical histone H2AK5ac modification, the ATAD2B bromodomain shows a two-fold higher affinity for the H2A.XK5ac mark (K*_D_* = 28.6 ± 2.0 µM), whereas the ATAD2 bromodomain demonstrates a 4-fold decrease in binding affinity for the same modification (K*_D_* = 160.3 ± 19.4 µM). Interestingly, while the ATAD2 and ATAD2B BRDs can interact with modified histone H2A.X, the ATAD2B BRD preferentially binds the modified histone H2A.XK5ac, while the ATAD2 BRD prefers the canonical H2A histone acetylated at K5. When we tested the binding activity of the ATAD2 and ATAD2B bromodomains with modifications on histone variant H2A.Z.1, the ATAD2B bromodomain binds with moderate affinity to the H2A.Z.1K4ac ligand (K*_D_* = 75.6 ± 4.1 µM). Interestingly, the ATAD2 bromodomain does not bind the mono- or the di-acetylated histone H2A.Z.1 ligands. Overall, our ITC data suggest that the ATAD2 bromodomain is more sensitive to changes in the H2A N-terminal amino acid sequence than the ATAD2B bromodomain, which exhibits a broader ligand binding specificity for acetylated lysine modifications on canonical and variant histones.

**Table 3:**
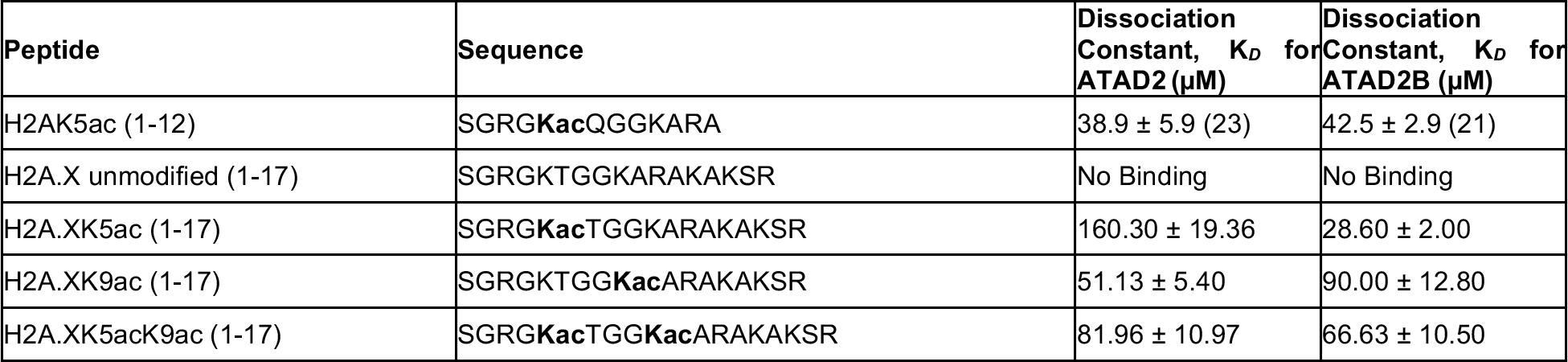
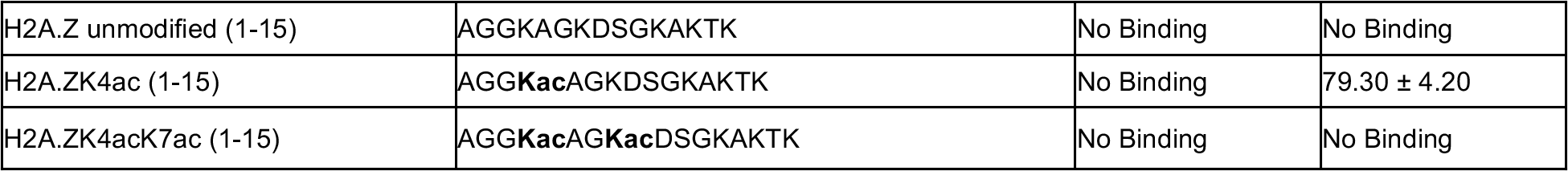
Histone variant H2A.X and H2A.Z N-terminus tail binding affinities to the ATAD2 and ATAD2B bromodomain-containing proteins. The apparent dissociation constants (*K_D_*) are given in micromolar. Amino acid mutations are highlighted in bold and underlined in the sequence.

## DISCUSSION

ATAD2 and its closely related paralog ATAD2B belong to sub-family IV of the human bromodomains (16). The bromodomains of these related proteins share a high degree of structural similarity and exhibit comparable ligand preferences for histone H4 acetylation modifications *in vitro* (20,21,39,40). Previous studies have found that the ATAD2 and ATAD2B bromodomains preferentially recognize di-acetylated histone H4K5acK12ac (19,21). However, a high-throughput combinatorial dCypher screen to identify post-translationally modified histone ligands of the ATAD2/B bromodomains indicated that the ATAD2B bromodomain was more promiscuous, binding 39 unique ligands, versus 11 bound by the ATAD2 bromodomain (21). Interestingly, the ATAD2 and ATAD2B bromodomains could bind to acetylated histone ligands containing distinctive arrangements of PTMs, including those that contained adjacent acetylation and methylation modifications (21). This result prompted us to further characterize the structural and functional differences between the ATAD2 and ATAD2B bromodomains to determine if they have distinctive histone-binding properties. These unique properties provide new insights into their unique cellular functions and may facilitate the development of specific agents that can target each protein by blocking their bromodomain acetyllysine binding activity.

ATAD2 has been widely studied in the context of cancer and is thought to be involved in chromatin remodeling activities associated with transcription, DNA damage repair, and cell cycle progression (19,20,46,47). However, the precise role of histone ligand recognition by the ATAD2 bromodomain in these various processes still needs further characterization. A recent study by Lazarchuk et al., showed that ATAD2 might play a role in DNA replication during the S phase of the cell cycle by protecting newly synthesized histones containing H4K5acK12ac modifications from the histone deacetylases HDAC1 and HDAC2 until they are incorporated into chromatin (46). Yet, the cellular function of ATAD2B is still unknown. Understanding the similarities and differences between the ATAD2 and ATAD2B bromodomain-histone PTM interactions may shed light on the unique functions of the ATAD2B protein.

In this study, we directly measured the binding affinities of the ATAD2/B bromodomains to histone H4 peptide ligands containing different combinations of PTMs. Our results demonstrated that the ATAD2 and ATAD2B bromodomains recognize distinct subsets of histone modifications. For example, the preferred mono-acetylated histone H4 ligand was H4K5ac for the ATAD2 bromodomain, whereas the ATAD2B bromodomain preferred H4K12ac. Structural analysis of the ATAD2/B BRDs in complex with the histone H4K12ac ligand (**Figure 2**) confirms that coordination of the acetyllysine moiety is consistent between the ATAD2 and ATAD2B BRDs (20). However, we found that there are distinct variations in how the histone backbone residues are coordinated by the bromodomains of ATAD2 and ATAD2B. For the ATAD2 BRD bound to H4K12ac (PDB ID 4QUT, **Figure 2D**), the histone ligand was coordinated mainly through water-mediated interactions between the protein and the histone peptide, except for histone Gly 9, which forms hydrogen bonds with D1066 and D1071 of the ATAD2 BRD (20). Alternatively, in ATAD2B BRD-H4K12ac structure (PDBID: 8EOQ), we observe an intricate network of direct hydrogen bonds between the histone ligand and the bromodomain binding pocket residues, which likely impart the higher selectivity of ATAD2B for the H4K12ac mark (**Figure 2C**). These distinctive differences in histone H4K12ac coordination by the ATAD2 and ATAD2B BRDs indicate they are utilizing unique binding modes.

Among the di-acetylated histone H4 ligands, both the ATAD2 and ATAD2B bromodomains preferred H4K5acK12ac (**Table 1**). This di-acetylated lysine combination is known to be highly enriched in newly synthesized histones in the nascent chromatin (47,48), and the full-length ATAD2 and ATAD2B proteins have also been shown to localize with newly synthesized chromatin (19,49). Thus, our ITC data demonstrating a strong affinity for the histone H4K5acK12ac ligand by the two bromodomains also supports their biological role in the regulation of nascent chromatin via interaction with di- acetylated H4 histones. Moreover, the di-acetylated H4K5acK12ac modification has been identified as an abundant proteoform in acute myeloid leukemia cells (50), which may recruit their activity in disease.

Within the chromatin, histones often contain multiple PTMs in various combinations (51–55). Hyper-acetylation of the histone H4 N-terminal tail has been associated with chromatin decompaction and transcriptional activation (56–59). Our ITC data demonstrated how multiple PTMs on the histone H4 N-terminus modulate the bromodomain activities of ATAD2 and ATAD2B. Within the multi-acetylated H4 peptides tested, the ATAD2 bromodomain strongly preferred the H4K8acK12acK16ac ligand, while the ATAD2B bromodomain selected the H4K5acK12acK16ac combination (**Table 1**). H4K8acK12acK16ac is one of the histone H4 hyper-acetylated proteoforms shown to increase dramatically upon HDAC inhibitor treatment of breast cancer cell lines SUM159 and MCF-7 (59).

Similarly, the histone H4R3me2a modification correlates with the transcriptional activation (24,60). However, in breast cancer patients, decreased abundance of the histone H4R3me2a mark was correlated with poor prognosis (61). Expression of the ATAD2 protein is up-regulated in breast cancer, and higher expression is correlated with poor patient prognosis (62). Our ITC data shows that the presence of bulky methyl groups on histone H4 inhibits the ATAD2 bromodomain activity, as demonstrated by the several-fold reduction in binding affinity of the histone H4R3me2aK5ac ligand when compared to the H4K5ac ligand (**Table 1**). Conversely, the ATAD2B bromodomain is more tolerant of the methylation modifications tested and showed no significant changes in binding affinity (**Table 1**).

Understanding the molecular mechanisms by which the ATAD2 and ATAD2B bromodomains recognize distinctive subsets of multivalent histones is essential for understanding their function as putative chromatin regulators. Histone H4 phosphorylation at Ser 1 (H4S1ph) is also associated with newly synthesized histones, and it is most abundant during the mitotic and S phases of the cell cycle (63). Phosphorylation of S1 on histone H4 is also induced following DNA damage and is possibly associated with DNA double-strand break repair (64). Our high-resolution crystal structures of ATAD2 and ATAD2B bromodomains in complex with the histone H4S1phK5ac ligand provide residue-specific information on how the addition of the negatively charged and bulky phosphoryl group at the N-terminus of histone H4 impacts the insertion of H4K5ac into the bromodomain pocket, resulting in altered bromodomain binding activity. In our previously solved structure of the ATAD2 BRD in complex with H4K5ac (PDB ID: 7M98) (23) without phosphorylation at Ser 1, the K5ac sidechain carbonyl forms a water-mediated hydrogen bond with the ATAD2 Y1021 residue (**Supp Figure S6**) (23). Moreover, the backbone carbonyl of Arg 3 of the H4K5ac ligand forms a direct hydrogen bond with the ATAD2 R1067 residue (23). These contacts are lost when Ser 1 is phosphorylated in the ATAD2 BRD - H4S1phK5ac complex, suggesting that phosphorylation of histone H4 at Ser 1 could function as a mechanism to prohibit recognition of transcriptionally active chromatin in the context of DNA damage.

Onco-histones have been described in various tumors where histone genes harbor mutations in regions encoding both the tail and core residues (41,42). Within histone H4, the most frequently observed mutations occur at the N-terminal tail. Our binding analyses demonstrated that mutations around the ‘Kac’ mark significantly impact the bromodomain activity (**Table 2**). Interestingly, mutations to Cysteine (e.g., S1C and R3C) increased the binding affinity for the ‘K5ac’ group by the ATAD2 BRD but reduced the binding affinity of the ATAD2B BRD. However, mutations to charged/polar residues (G4D and G4S) abolished binding to the adjacent K5ac mark by both bromodomains. Based on our ATAD2 and ATAD2B BRD structures in complex with the histone H4S1CK5ac ligand, the altered binding affinities for these mutated onco-histones likely result from changes in the polar and hydrophobic contacts between the histone ligand and bromodomain. Our structure-function analysis suggests that onco-histone mutations of Gly 4 to an Asp or Ser residue would cause steric clashes from unfavorable orientations of their bulky sidechains within the bromodomain binding pocket, prohibiting histone recognition. Thus, onco-histone mutations have the potential to play a significant role in inhibiting the activity of bromodomain-containing proteins in cancer.

Like the canonical histones H3, H4, H2A, and H2B, histone protein variants can also have a variety of post-translational modifications (44,65). However, due to variations in their amino-acid sequence, the histone variants create unique binding sites for proteins that read post-translational modifications. Despite the importance of histone variants in essential cellular processes, including the regulation of transcription, DNA repair, and mitosis (66,67), the interaction of chromatin ‘readers’ with modified histone variants has not been actively studied. This study focused on the interaction of acetylated histone H2A.X and H2A.Z variants with the ATAD2 and ATAD2B bromodomains and demonstrated how changes in the histone backbone sequence modulate the recognition of the ‘Kac’ PTM. The ATAD2 bromodomain was found to be more selective, and changes in the histone sequence surrounding the ‘Kac’ modification resulted in much weaker binding affinities for ATAD2 with PTM histone variants. Notably, the ATAD2B bromodomain is preferentially bound to the H2A.XK5ac modification over the canonical histone H2AK5ac mark, and it was also able to interact with the acetylated histone variant H2A.ZK4ac, while the ATAD2 bromodomain was not. Our results align with the cell-cycle expression pattern of ATAD2, which is predominantly expressed during the S phase when newly synthesized histones are most abundant (19). In contrast, the histone variants are expressed throughout the cell cycle (68,69). Notably, insertion and acetylation of H2A.Z at K4, K7, and K11 are associated with actively transcribed chromatin at promoter regions, where it functions to destabilize DNA wrapping and opening the nucleosome stacks (35,70–72). Recently Kim JJ et al. reported a non-homologous end-joining DNA repair phenotype associated with ATAD2B in a bromodomain-containing protein-wide siRNA screen (73). This result was not followed up, but the data suggest that, unlike its ATAD2 paralog, ATAD2B may have activity outside of the S phase. The histone H2A.X variant is strongly associated with DNA damage repair through the crucially important S139ph signaling event, also known as γ-H2A.X. The abundance of H2A.X relative to total cellular H2A varies greatly, from 2% to 36%, in the sparse reports available (74,75). Histone acetyltransferase TIP60 also acetylates histone H2A.X at the K5 position after DNA damage, creating H2A.XK5ac, which accelerates histone exchange at sites of DNA double-strand breaks (DSB) (25,76). Moreover, the turnover of NBS1, a primary sensor of double strand breaks at DNA damage sites, depends on the acetylation state of histone H2A.X(76). Our ITC binding data demonstrates that the ATAD2 and ATAD2B bromodomains have the ability to interact with N-terminally acetylated H2A.X, which is consistent with their potential roles in DNA damage repair (73,77).

The interplay between diverse histone PTMs forms the ‘histone code,’ which drives the recruitment of reader proteins to the chromatin and ultimately dictates specific cellular outcomes. Our current study highlights the importance of studying crosstalk between various histone post-translational modifications to understand how they modulate bromodomain-containing protein binding activity. Adjacent histone PTMs can either promote or inhibit the affinity of bromodomains with chromatin, which, in turn, regulates essential cellular processes such as gene transcription (**Figure 5**). Future investigations on how individual and combinatorial histone modifications direct the localization of ATAD2 and ATAD2B to specific regulatory sites along the genome are needed to understand how these complex epigenetic signals created by the ‘histone code’ drives gene regulation in specific biological contexts.

**Figure 5:**
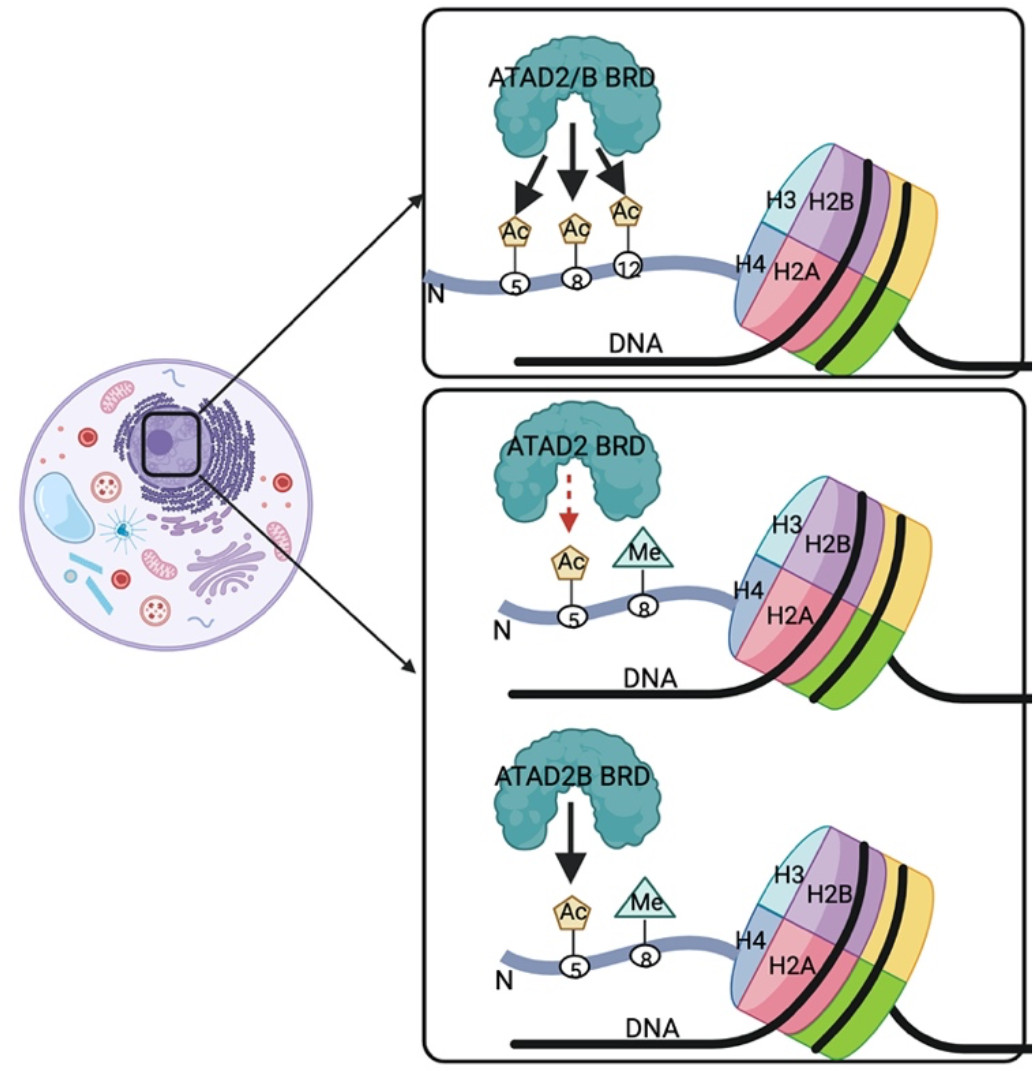
Multiple modifications on individual histone tails of the nucleosome modulate the ATAD2/B bromodomain activity. In the cell, ATAD2/B is localized at the chromatin, which is enriched with histone PTMs. These modifications influence the ATAD2/B bromodomain activity. Adjacent methyl groups negatively impact the ATAD2 bromodomain’s ability to recognize the ‘Kac’ mark but do not significantly affect the ATAD2B bromodomain activity.

In summary, a comparative evolutionary analysis of ATAD2 and ATAD2B highlights their high conservation as paralogues, characterized by gene duplication and subsequent evolution in chordates to acquire distinct bromodomain binding activities. Our findings strongly suggest the likelihood of non-redundant functions for the ATAD2 and ATAD2B bromodomain-containing proteins and emphasize the important role of their bromodomains in directing their unique biological functions. Bromodomain inhibitors have emerged as a primary research focus, and given the association of ATAD2 as a key player in cancer, targeting the specific bromodomain activities of ATAD2 and ATAD2B could be vital to developing potent therapeutic interventions.

## Author Contributions

Conceptualization, M.P., K.L.M., S.E.F., and K.C.G.; investigation and validation, M.P., K.L.M., B.W.B., C.M., I.A.K., F.M.J., K.M.B., S.P.B., S.C., J.C.N., N.L.Y., V.J., S.E.F., and K.C.G.; writing, M.P., reviewing, and editing M.P., K.L.M., F.M.J., K.M.Q., N.L.Y., V.J., S.E.F., and K.C.G.; supervision, S.E.F., N.L.Y., and K.C.G.; funding acquisition, K.C.G., and S.E.F. All authors have read and agreed to the final version of the manuscript.

## DATA AVAILABILITY

The atomic coordinates and structure factors for the ATAD2 and ATAD2B BRD in complex with histone ligands - H4K12ac (4-17), H4S1phK5ac (1-15), and H4S1CK5ac (1-15) have been deposited in PDB and will be made available upon publication. The mass spectrometry data will be available through the ProteomeXchange consortium upon publication.

## ACCESSION NUMBERS

The atomic coordinates and structure factors for the ATAD2 BRD in complex with H4S1phK5ac (1-15), H4S1CK5ac (1-15), and ATAD2B BRD in complex with histones H4K12ac (4-17), H4S1phK5ac (1-15), and H4S1CK5ac (1-15) have been deposited with the Protein Data Bank under the accession numbers – 8SDQ, 8SDO, 8EOQ, 8ESJ, and 8SDX respectively.

