## Supplementary information for "Impact of combinatorial histone modifications on acetyllysine recognition by the ATAD2 and ATAD2B bromodomains"

*M.P. and K.L.M. share first authorship

**Table of Contents**

Supp. Fig. S1: Histone H4 peptides with combinatorial PTMs binding to the ATAD2 bromodomain. . ....................................................................................................3

Supp. Fig. S2: Histone H4 peptides with combinatorial PTMs binding to the ATAD2B bromodomain. . ....................................................................................................4

Supp. Fig. S5: Histone H2A.X and H2A.Z variant peptides with combinatorial PTMs binding to the ATAD2/B bromodomain. . .....................................................................10

Supp. Fig. S6: Coordination of the histone H4 ligand with K5ac by the ATAD2 bromodomain…………………………………………………………………………..……... 11

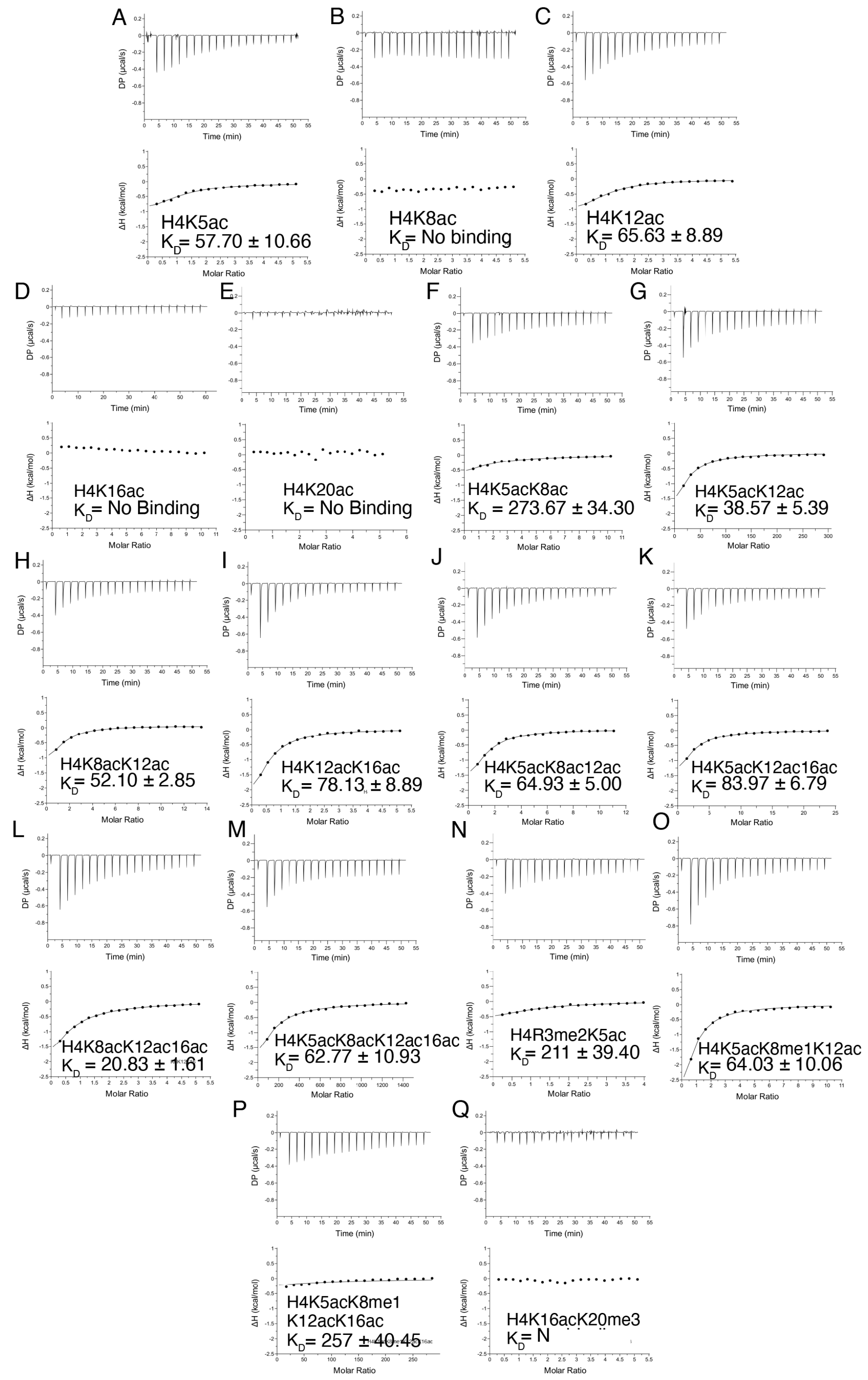

**Supp. Fig. S1: Histone H4 peptides with combinatorial PTMs binding to the ATAD2 bromodomain.** A-Q) Exothermic enthalpy ITC curves for the H4 peptides with various combinations of acetylation, methylation, and phosphorylation. The calculated binding constant is given in $\mu$M as $K_D$ for each peptide. All peptides are 1-24 amino acids in length.

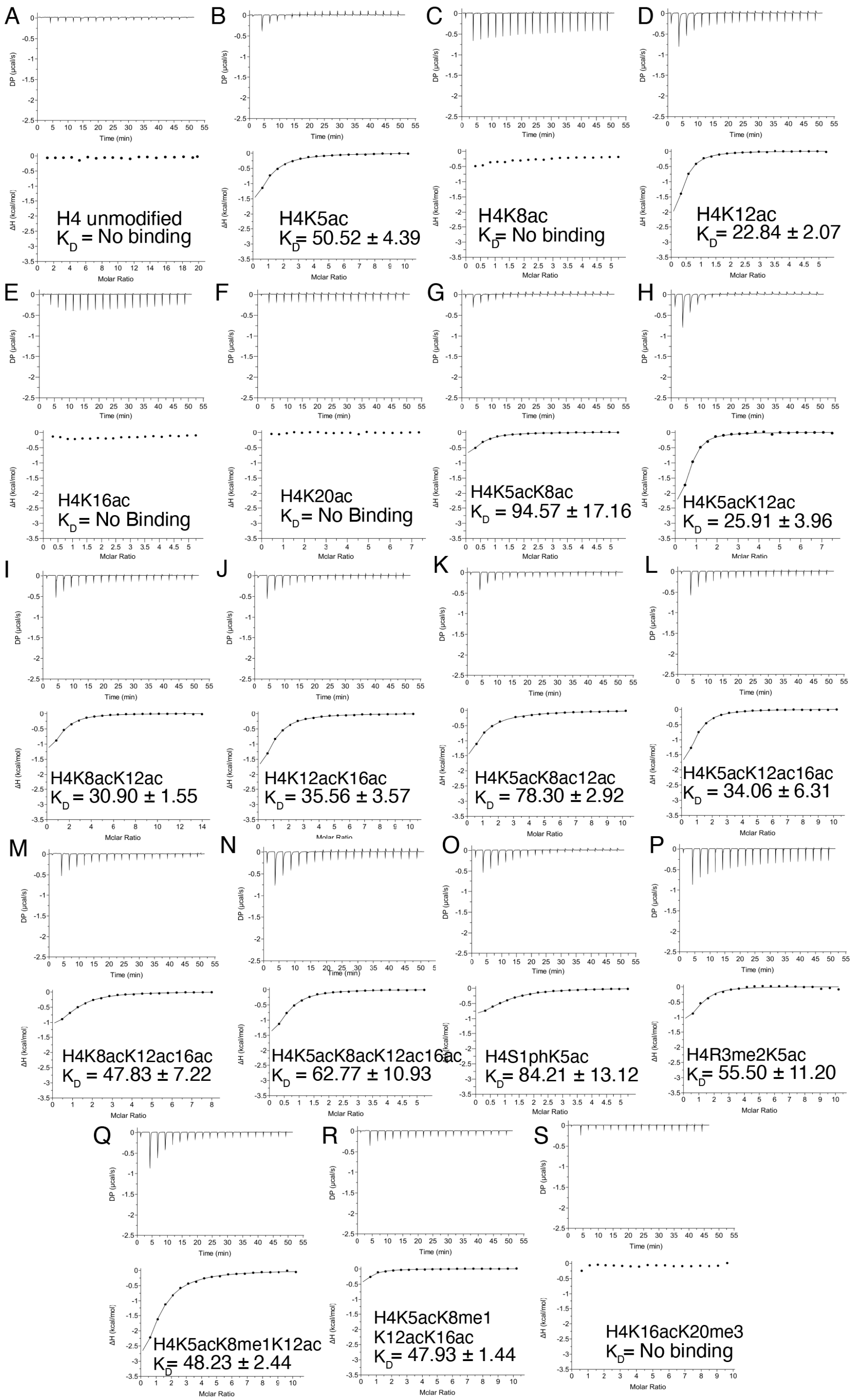

**Supp. Fig. S2: Histone H4 peptides with combinatorial PTMs binding to the ATAD2B bromodomain.** A-S) Exothermic enthalpy ITC curves for the H4 peptides with various combinations of acetylation, methylation, and phosphorylation. The calculated binding constant is given in μM as $K_D$ for each peptide. All peptides are 1-24 amino acids in length.

**Supp. Table S1: Summary of the data collection and refinement statistics for the crystal structure of ATAD2B BRD in complex with H4K12ac** (PDB ID: 8EOQ).

| | |
|---|---|
| Wavelength | 1 |
| Resolution range | 27.56 - 2.0 (2.072 - 2.0) |
| Space group | P 41 21 2 |
| Unit cell | 55.119 55.119 140.657 90 90 90 |
| Total reflections | 206211 (19202) |
| Unique reflections | 15396 (1490) |
| Multiplicity | 13.4 (12.9) |
| Completeness (%) | 99.84 (99.67) |
| Mean I/sigma(I) | 22.67 (1.68) |
| Wilson B-factor | 34.01 |
| R-merge | 0.1008 (1.635) |
| R-meas | 0.1048 (1.702) |
| R-pim | 0.02856 (0.471) |
| $CC_{1/2}$ | 0.999 (0.755) |
| CC* | 1 (0.928) |
| Reflections used in refinement | 15385 (1489) |
| Reflections used for R-free | 729 (65) |
| R-work | 0.1979 (0.2587) |
| R-free | 0.2191 (0.2728) |
| CC(work) | 0.968 (0.865) |
| CC(free) | 0.949 (0.830) |
| Number of non-hydrogen atoms | 1332 |
| macromolecules | 1232 |
| solvent | 100 |
| Protein residues | 151 |
| RMS(bonds) | 0.011 |
| RMS(angles) | 1.26 |
| Ramachandran favored (%) | 99.30 |
| Ramachandran allowed (%) | 0.70 |
| Ramachandran outliers (%) | 0.00 |
| Rotamer outliers (%) | 0.00 |
| Clashscore | 3.20 |
| Average B-factor | 41.69 |
| macromolecules | 41.64 |
| solvent | 42.26 |

Statistics for the highest-resolution shell are shown in parentheses.

**Supp. Table S2: Summary of data collection and refinement statistics for the crystal structure of ATAD2 BRD in complex with H4S1phK5ac** (PDBID: 8SDQ).

| | |
|---|---|
| Wavelength | 1.072 |
| Resolution range | 44.34 - 1.85 (1.916 - 1.85) |
| Space group | 44.34 - 1.85 (1.916 - 1.85) |
| Unit cell | 52.893 52.893 162.602 90 90 90 |
| Total reflections | 283313 (25897) |
| Unique reflections | 20634 (1659) |
| Multiplicity | 13.7 (12.8) |
| Completeness (%) | 97.37 (82.12) |
| Mean I/sigma(I) | 26.44 (3.15) |
| Wilson B-factor | 23.67 |

| | |
|---|---|
| R-merge | 0.08953 (0.9309) |
| R-meas | 0.09301 (0.9695) |
| R-pim | 0.02498 (0.2679) |
| $CC_{1/2}$ | 1 (0.913) |
| CC* | 1 (0.977) |
| Reflections used in refinement | 20095 (1658) |
| Reflections used for R-free | 996 (74) |
| R-work | 0.2047 (0.3835) |
| R-free | 0.2379 (0.4538) |
| CC(work) | 0.959 (0.608) |
| CC(free) | 0.956 (0.256) |
| Number of non-hydrogen atoms | 1396 |
| macromolecules | 1186 |
| solvent | 210 |
| Protein residues | 140 |
| RMS(bonds) | 0.011 |
| RMS(angles) | 1.10 |
| Ramachandran favored (%) | 97.73 |
| Ramachandran allowed (%) | 2.27 |
| Ramachandran outliers (%) | 0.00 |
| Rotamer outliers (%) | 0.00 |
| Clashscore | 3.81 |
| Average B-factor | 34.24 |
| macromolecules | 32.79 |
| solvent | 42.40 |
| Statistics for the highest-resolution shell are shown in parentheses | |

**Supp. Table S3: Summary of data collection and refinement statistics for the crystal structure of ATAD2B BRD in complex with H4S1phK5ac** (PDBID: 8ESJ).

| | |
|---|---|
| Wavelength | 1.072 |
| Resolution range | 31.37 - 1.4 (1.45 - 1.4) |
| Space group | P 21 21 21 |
| Unit cell | 41.08 48.58 83.32 90 90 90 |
| Total reflections | 397691 (37316) |
| Unique reflections | 33447 (3285) |
| Multiplicity | 11.9 (11.4) |
| Completeness (%) | 99.51 (99.57) |
| Mean I/sigma(I) | 28.93 (2.50) |
| Wilson B-factor | 18.27 |
| R-merge | 0.05652 (1.989) |
| R-meas | 0.05916 (2.082) |
| R-pim | 0.01714 (0.6077) |
| $CC_{1/2}$ | 1 (0.873) |
| CC* | 1 (0.965) |
| Reflections used in refinement | 33431 (3275) |
| Reflections used for R-free | 1678 (154) |
| R-work | 0.1784 (0.2520) |
| R-free | 0.2023 (0.2784) |
| CC(work) | 0.968 (0.895) |
| CC(free) | 0.964 (0.832) |
| Number of non-hydrogen atoms | 1399 |
| macromolecules | 1131 |

| | |
|---|---|
| solvent | 268 |
| Protein residues | 138 |
| RMS(bonds) | 0.011 |
| RMS(angles) | 1.04 |
| Ramachandran favored (%) | 97.71 |
| Ramachandran allowed (%) | 2.29 |
| Ramachandran outliers (%) | 0.00 |
| Rotamer outliers (%) | 0.00 |
| Clashscore | 3.10 |
| Average B-factor | 29.77 |
| macromolecules | 27.88 |
| solvent | 37.77 |
| Statistics for the highest-resolution shell are shown in parentheses | |

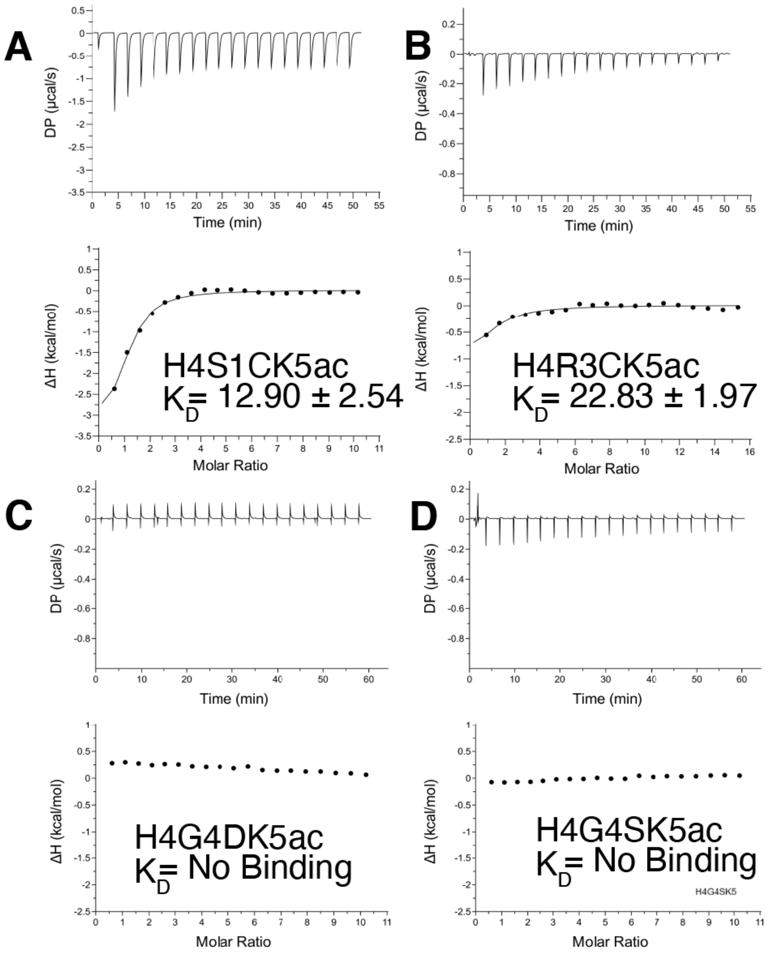

**Supp. Fig. S3: Binding of 'oncohistone' H4 peptides to the ATAD2 bromodomain.** A-D) ITC enthalpy plots for the binding of the ATAD2 BRD with mutant mono-acetylated histone H4 ligands (residues 1-15) as indicated. The calculated binding constant ($K_D$) for each peptide is shown in their respective plots.

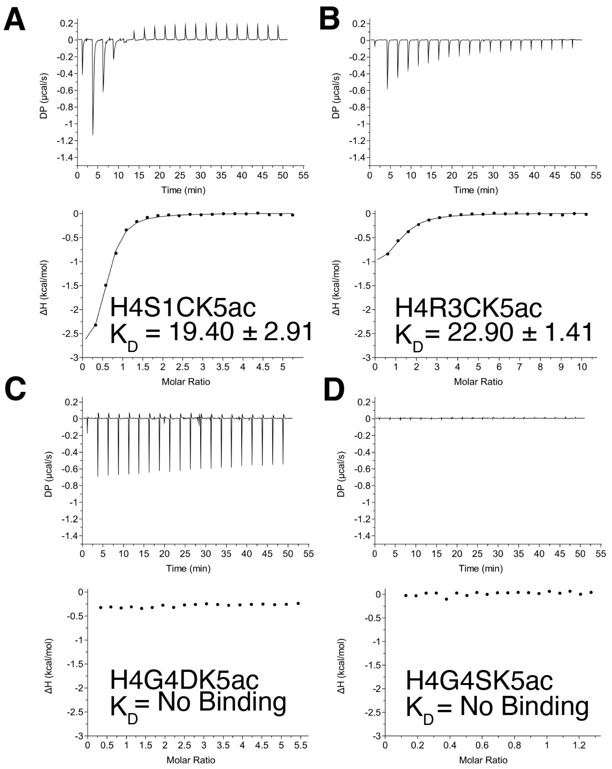

**Supp. Fig. S4: Binding of 'oncohistone' H4 peptides to the ATAD2B bromodomain.** A-D) ITC enthalpy plots for the binding of the ATAD2B BRD with mutant mono-acetylated histone H4 ligands (residues 1-15) as indicated. The calculated binding constant ($K_D$) for each peptide is shown in their respective plots.

**Supp. Table S4: Summary of data collection and refinement statistics for the crystal structure of ATAD2 BRD in complex with H4S1CK5ac** (PDBID: 8SDO).

| | |
|---|---|
| Wavelength | 1.072 |
| Resolution range | 27.46 - 2.01 (2.082 - 2.01) |
| Space group | P 43 2 2 |
| Unit cell | 53.941 53.941 159.522 90 90 90 |
| Total reflections | 220898 (21350) |
| Unique reflections | 16480 (1529) |
| Multiplicity | 13.4 (13.4) |
| Completeness (%) | 99.51 (96.10) |
| Mean I/sigma(I) | 18.91 (0.65) |
| Wilson B-factor | 43.36 |
| R-merge | 0.1117 (4.016) |
| R-meas | 0.1162 (4.175) |
| R-pim | 0.0316 (1.133) |
| $CC_{1/2}$ | 0.999 (0.374) |
| CC* | 1 (0.738) |
| Reflections used in refinement | 16410 (1529) |
| Reflections used for R-free | 762 (67) |
| R-work | 0.2009 (0.3077) |
| R-free | 0.2306 (0.3278) |
| CC(work) | 0.966 (0.665) |
| CC(free) | 0.966 (0.711) |
| Number of non-hydrogen atoms | 1309 |
| macromolecules | 1141 |
| ligands | 0 |
| solvent | 168 |
| Protein residues | 136 |
| RMS(bonds) | 0.007 |
| RMS(angles) | 0.95 |
| Ramachandran favored (%) | 98.46 |
| Ramachandran allowed (%) | 1.54 |
| Ramachandran outliers (%) | 0.00 |
| Rotamer outliers (%) | 1.59 |
| Clashscore | 3.97 |
| Average B-factor | 56.87 |
| macromolecules | 55.63 |
| solvent | 65.29 |

Statistics for the highest-resolution shell are shown in parentheses.

**Supp. Table S5: Summary of data collection and refinement statistics for the crystal structure of ATAD2B BRD in complex with H4S1CK5ac** (PDBID: 7TZQ).

| | |
|---|---|
| Wavelength | 1.00 |
| Resolution range | 33.52 - 2.79 (2.89 - 2.79) |
| Space group | P 1 21 1 |
| Unit cell | 47.687 79.629 64.348 90 105.141 90 |
| Total reflections | 42819 (3969) |
| Unique reflections | 11594 (1060) |
| Multiplicity | 3.7 (3.5) |
| Completeness (%) | 98.41 (92.81) |
| Mean I/sigma(I) | 8.41 (1.21) |
| Wilson B-factor | 65.79 |
| R-merge | 0.1152 (1.246) |
| R-meas | 0.1351 (1.474) |
| R-pim | 0.06988 (0.782) |
| $CC_{1/2}$ | 0.994 (0.524) |
| CC* | 0.998 (0.829) |
| Reflections used in refinement | 11481 (1059) |
| Reflections used for R-free | 598 (53) |

| | |
|---|---|
| R-work | 0.2208 (0.3775) |
| R-free | 0.2563 (0.4409) |
| CC(work) | 0.960 (0.678) |
| CC(free) | 0.968 (0.503) |
| Number of non-hydrogen atoms | 2342 |
| macromolecules | 2311 |
| ligands | 5 |
| solvent | 26 |
| Protein residues | 284 |
| RMS(bonds) | 0.011 |
| RMS(angles) | 1.41 |
| Ramachandran favored (%) | 93.70 |
| Ramachandran allowed (%) | 5.93 |
| Ramachandran outliers (%) | 0.37 |
| Rotamer outliers (%) | 1.56 |
| Clashscore | 15.65 |
| Average B-factor | 73.62 |
| macromolecules | 73.77 |
| ligands | 66.62 |
| solvent | 61.46 |

Statistics for the highest-resolution shell are shown in parentheses.

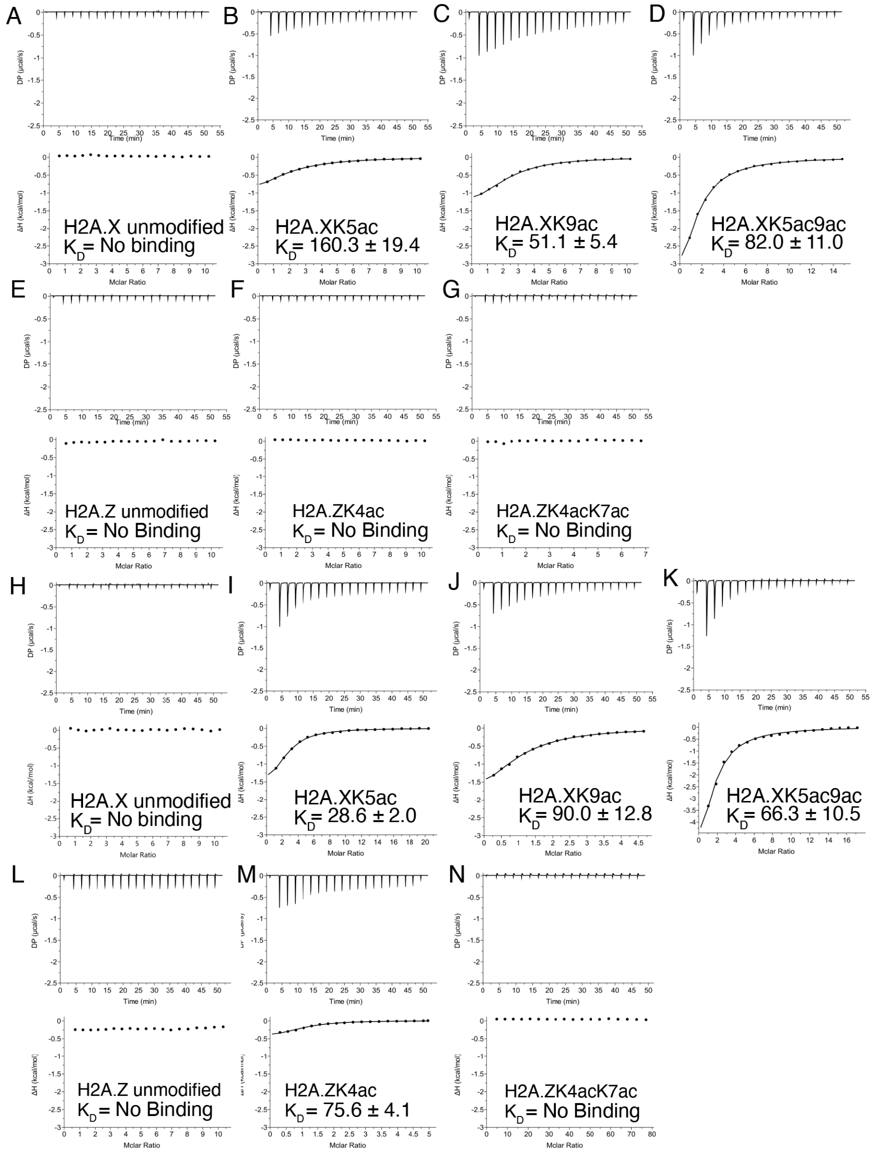

**Supp. Fig. S5: Histone H2A.X and H2A.Z variant peptides with combinatorial PTMs binding to the ATAD2/B bromodomain.** A-G) Exothermic enthalpy ITC curves for the H2A.X and H2A.Z histone variant peptides with mono- and di-acetylation marks binding to ATAD2 bromodomain. H-N) Exothermic enthalpy ITC curves for the H2A.X and H2A.Z histone variant peptides with mono- and di-acetylation marks binding to ATAD2B bromodomain. The calculated binding constant is given in μM as $K_D$ for each peptide.

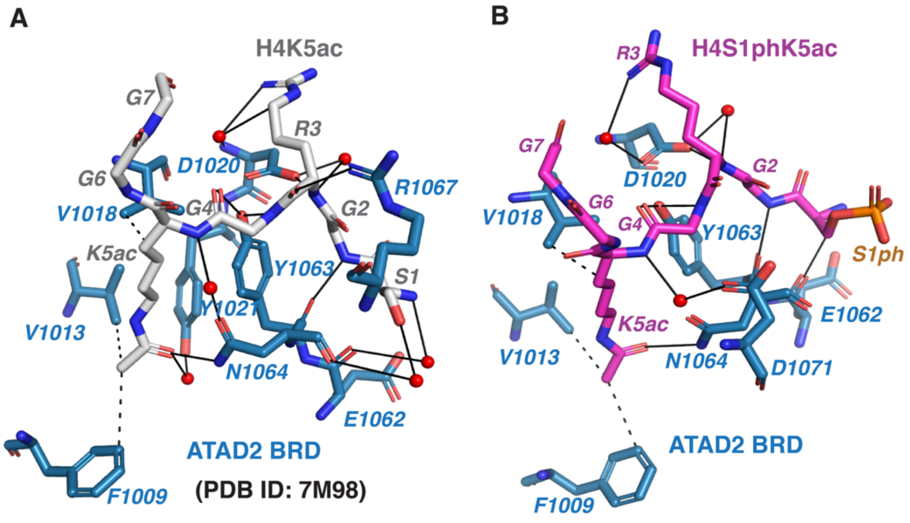

**Figure S6: Coordination of the histone H4 ligand with K5ac by the ATAD2 bromodomain.** (A) Coordination of the H4K5ac (residues 1-10) by the ATAD2 BRD observed in the previously solved crystal structure PDB ID: 7M98(1). ATAD2 BRD residues are in blue, while the histone H4K5ac ligand residues are in grey. (B) Coordination of the H4S1phK5ac ligand (residues 1-7) by the ATAD2 BRD. ATAD2 BRD residues are displayed in dark blue, while the H4S1phK5ac ligand residues are shown in magenta. Solid black lines represent hydrogen bonds, while dashed black lines show hydrophobic interactions. Water molecules are shown in red. All figures were made using the PyMOL Molecular Graphics system using version 2.3, Schrödinger, LLC, and all contacts were determined using the PLIP program (2).
